# Disruption of a side portal pathway permits closed-state inactivation by BK β subunit N-termini

**DOI:** 10.1101/2025.03.09.642150

**Authors:** Yu Zhou, Xiao-Ming Xia, Christopher J. Lingle

## Abstract

Cytosolic N-termini of several BK channel β regulatory subunits mediate rapid inactivation. However, in contrast to Kv channels, inactivation does not occur via a simple, open channel block mechanism, but involves two steps, an association step in which ion permeation is maintained (O*), then followed by inactivation (I). To produce inactivation, BK β subunit N-termini enter the central cavity through a lateral entry pathway (“side portal”) separating the transmembrane pore-gate-domain and cytosolic gating ring. Comparison of BK conformations reveals an aqueous pathway into the central cavity in the open structure, while in the closed structure three sequential basic residues (R_329_K_330_K_331_) in S6 occlude central cavity access. We probed the impact of mutations of the RKK motif (RKK3Q, RKK3E, and RKK3V) on inactivation mediated by the β3a N-terminus. All three RKK-mutated constructs differentially reduce depolarization-activated outward current, prolong β3a-mediated tail current upon repolarization, and produce a persistent inward current at potentials down to −240 mV. With depolarization channels are driven into O*-I inactivated states and, upon repolarization, slow tails and persistent inward currents reflect slow changes in O*-I occupancy. However, evaluation of closed state occupancy prior to depolarization and at the end of slow tails reveals that some fraction of closed states at negative potentials corresponds to resting closed states in voltage-independent equilibrium with N-terminal-occluded closed-states. Thus, disruption of the RKK triplet both stabilizes the β3a-N-terminus in its position of inactivation and permits access of that N-terminus to its blocking position in closed states.

**Summary:** The role of BK S6 residues R329K330K331 and E321/E324 in β subunit-mediated inactivation is probed. WT R329K330K331 hinders inactivation in closed states, while RKK mutations stabilize inactivated states even under conditions where channels are otherwise closed. E321/E324 mutations do not permit closed-state inactivation.

## Introduction

Ca^2+^ and voltage-activated, large conductance BK-type K^+^ channels undergo a fast inactivation process that arises from N-terminal peptide segments of regulatory β2 (Wallner et al., 1999; Xia et al., 1999) and β3a, β3b, and β3c subunits (Uebele et al., 2000; Xia et al., 2000; Zeng et al., 2007). Although this process qualitatively shares similarities to *Shaker* fast inactivation (Hoshi et al., 1990; Zagotta et al., 1990), inactivation mediated by BK β subunits exhibits features inconsistent with simple, open-channel block. Cytosolic channel blockers do not compete with β subunit mediated inactivation (Solaro et al., 1997; Xia et al., 2000), although TEA has been shown to compete with an exogenously applied N-terminal peptide blocker (Wallner et al., 1999). The presence of time-dependent changes in instantaneous IV curves during development of inactivation support the existence of two categories of open state (Lingle et al., 2001; Benzinger et al., 2006), one that closes normally and one in rapid equilibrium with inactivated states. Finally, tail current behavior during recovery from inactivation is inconsistent with the predictions of recovery from simple open-channel block (Solaro et al., 1997; Zeng et al., 2007; Gonzalez-Perez et al., 2012). All three of these features of BK inactivation have been explained by a two-step inactivation process which in its simplest form is given by:

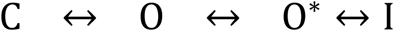

The key point encapsulated in such a scheme is that a rate-limiting transition to a pre-inactivated open state (O*) precedes a rapid transition to a non-conducting inactivated state. For the β3a N-terminus, repeated cycling between I and O* explains the prolonged tail of current following repolarization after channels are inactivated at depolarized potentials. For β3a, this mechanism is of potential physiological significance, distinguishing it from simple channel block (C←→O←→I), since it results in a much larger net current flux (Zeng et al., 2007; Gonzalez-Perez et al., 2012) than would be predicted from simple open channel block (Demo and Yellen, 1991; Raman, 2023).

An additional intriguing difference between BK and *Shaker* inactivation concerns stereospecificity in the channel blocking effects of isolated N-terminal peptides. *Shaker*-inactivation-removed (*Shaker*-IR) channels are inhibited essentially indistinguishably between D- and L-20 mer N-terminal *Shaker* peptides (Gonzalez-Perez et al., 2012). In contrast, the 20-mer L-peptide of the BK β3a N-terminus recapitulates all the features of the two-step inactivation process, while a D-β3a peptide exhibits features of simple channel block characteristic of *Shaker* (Gonzalez-Perez et al., 2012). This suggests that BK inactivation involves a stereospecific binding step that precedes inactivation. Yet, a challenge for understanding BK inactivation has been that, whereas *Shaker*-type N-terminal inactivation can be easily conceptualized by the simple block, pore-occlusion model (Hoshi et al., 1990; Zagotta et al., 1990; Choi et al., 1991; Demo and Yellen, 1991; Aldrich, 2001), for BK inactivation the structural correlates that may underlie the postulated two-steps in the inactivation process are lacking. Furthermore, the quite different β2 and β3a tail current behaviors have not been fully explained (Zeng et al., 2007; Gonzalez-Perez et al., 2012).

BK β subunit N-termini access their positions of inactivation, presumably within the central cavity, via passage through side portals that connect the membrane associated pore-gate-domain (PGD) to the cytosolic domain (CTD) containing the gating rings (Zhang et al., 2006; Zhang et al., 2009; Agarwal et al., 2025). Reasoning that some of the unique features of BK β subunit inactivation might reflect interactions of the N-termini with the access pathway, here we have taken advantage of current BK structural information (Hite et al., 2017; Tao et al., 2017; Tao and MacKinnon, 2019) to probe a potential pathway by which β subunit N-termini might reach the BK central cavity. We find that, in the available structural model of the unliganded apo BK channel, there is no obvious pathway that would accommodate movement of N-terminal segments to the central cavity, while in the liganded model a pathway can be identified. The key structural difference between the two conformations is that a triplet of basic residues, R_329_K_330_K_331_ (RKK), is positioned to hinder β subunit N-terminal passage into the central cavity in the closed state. The RKK triplet has received extensive attention in previous work with results suggesting that the S6-RCK1 linker segment that contains the RKK triplet is a critical structural determinant of multiple aspects of gating. This includes stabilization of the closed BK conformation by interaction between RKK and two acid residues (E321/E324) from an adjacent subunit (Tian et al., 2015), participation in sensitivity to PIP2 (Vaithianathan et al., 2008; Tian et al., 2015), coupling of the cytosolic gating ring (CTD) to pore-gate domain (PGD) function (Niu et al., 2004), and coupling of voltage-sensor activation to channel opening (Sun and Horrigan, 2022). Here we utilize macroscopic and single channel evaluation of RKK (and E321/E324) mutations to address their impact on inactivation mediated by β3a and β2 subunits. The results suggest that each of the RKK triplet mutations differentially stabilize the lifetime of channels in inactivated states. Furthermore, contrary to the linear formulation presented above, disruption of the RKK triplets also permits direct inactivation from resting closed states. In contrast, mutations of E321/E324 predominantly reduce the stability of the inactivated states. The results support the view that disruption of the RKK motif creates a pathway in closed channels that allows N-terminal inactivation motifs to reach a position of inactivation. Our estimates suggest that the equilibrium constants for binding of the N-terminus to either open or closed states in a given RKK mutant channel are not too different.

## Materials and Methods

### Molecular Biology and Channel Expression

BK channels were expressed in stage IV Xenopus oocytes by cRNA injection of the pore forming mouse Slo1 (mSlo1) α-subunit alone or with hβ2 or D20Am (Gonzalez-Perez et al., 2012). The cRNAs of α and β subunits were injected at 1:4 ratio by weight to ensure molar excess of β subunits. Three constructs of the mSlo1 α subunit were generated in which the triplet of residues, R_329_K_330_K_331_, immediately following the S6 pore-lining helix, were mutated. Construct RKK/3Q corresponds to replacement of RRK with three glutamine residues, RKK/3E corresponds to replacement with glutamate residues, and RKK/3V corresponds to replacement by valine. A second set of mutant constructs focused on two acidic residues near the end of the S6 helix, E321 and E324. Here we utilized EE/QQ, EE/DD, EE/GG, EE/WW, and EE/KK constructs. All mutants were generated by site-directed mutagenesis. The specific DNA primers were purchased from Integrated DNA Technologies (Coralville, IA) and Pfu DNA polymerase was applied to perform 18 PCR cycles (96°C, 30’’; 52°C, 30”; 69°C, 18’). After 2 hours DpnI digestion, the reactions were used to transform E.coli DH10B electrically (Invitrogen, Carlsbad, CA), which cells were then plated on LB plates containing 70 µg/mL carbenicillin. The resulting single colonies were grown out for plasmid DNA isolations. The intended mutants were identified and verified through Sanger sequencings (Azenta Life Sciences, Plainfield NJ). To synthesize cRNA, the pCap vector templates were linearized with MluI, purified, and cRNAs were synthesized with T7 RNA polymerase (Invitrogen).

### Measurement and Visualization of BK Aqueous Accessible Volume

The aqueous accessible volumes in hSlo1 structures were evaluated using the program PrinCCes (Czirjak, 2015) with a probe of 8Å diameter. The structural images were prepared using UCSF ChimeraX (Goddard et al., 2018).

### Electrophysiology and Data Analysis

All currents were recorded in the inside-out patch configuration using an Axopatch 200B amplifier (Molecular Devices, Sunnyvale, CA). Currents were low-pass filtered at 10 kHz and digitized at 100 kHz. During seal formation, an oocyte was bathed in standard frog Ringer (in mM): 115 NaCl, 2.5 KCl, 1.8 CaCl_2_, 10 HEPES, at pH 7.4. Following patch excision, the pipette tip was moved into flowing test solutions of defined Ca^2+^ concentrations. The pipette was filled with extracellular solution containing (in mM): 140 K-methanesulfonate, 20 KOH, 10 HEPES, 2 MgCl_2_, at pH 7.0. The intracellular solutions included (in mM): 140 K-methanesulfonate, 20 KOH, 10 HEPES at pH 7. 0. 5 mM HEDTA was used for 10 μM Ca^2+^ and 5 mM EGTA for 0 mM Ca^2+^ solutions.

For single channel patches, currents were sampled at 50 kHz and filtered at 5 kHz. Leak and capacity transients were subtracted by using blank traces in the same recording. For cases of BK channels that were open at the end of the depolarization to +140 mV, only rarely would a fully resolved deactivating BK channel be expected to be observed. BK channel deactivation at −160 mV typically occurs with a time constant of about 100 μs (Horrigan and Aldrich, 2002), which predicts that less than 5% of all channel closing events will be longer than 300 μs. For the 36 of 860 cases in which WT+D20Am channels were open at the time of repolarization, only 1 or 2 deactivation events longer than 300 μs would be expected.

The trypsin digestion procedure was as described previously (Zhang et al., 2006; Zhang et al., 2009). In short, trypsin was applied for timed durations in 0 [Ca^2+^]_in_ while the patch was held at −70 mV. Digestion was then monitored following return to a 10 μM [Ca^2+^]_in_ solution lacking trypsin using currents evoked by test steps to +160 mV after a preconditioning step to −160 mV. Each patch was washed in enzyme-free 0 [Ca^2+^]_in_ solution for at least 5 s before and after each test current recording.

Solutions at the pipette tip of excised inside-out patches were changed with an SF-77B fast perfusion stepper system (Warner Instruments) controlled by pClamp (Molecular Devices, San Jose, CA). Experiments were performed at room temperature (∼22-25 °C). All chemicals were purchased from Sigma-Aldrich (St. Louis, MO).

The G-V relationship of BK channels was determined from tail currents measured 150 μs after repolarization to −120 mV. G-V curves were fit by a Boltzmann function of the form:

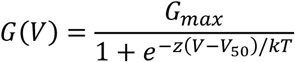

in which G_max_ is maximal conductance, *z* is apparent voltage-dependence in units of elementary charge, V_50_ is the voltage of half-maximal activation, and k and T have their usual physical meanings.

Data were analyzed using OriginPro 7.5 (OriginLab Corporation), Clampfit (Molecular Devices, San Jose, CA), GraphPad Prism 10.4.0 (GraphPad, San Diego, CA) and programs developed in this laboratory. Error bars in the figures represent standard deviations (SD). Each measurement is averaged from at least 4 experiments with number of patches given in figure legends. Specific statistical tests are given when used. For comparisons of single channel events among constructs, Fisher’s exact test compared 2×2 contingency matrices was used. To correct for multiple comparisons, a Bonferroni correction was used, such that for any comparison of events in two constructs, the P-value reported by the Fisher test was corrected to P times the number of different comparisons.

### Analysis of peak current reduction mediated by D20Am expressed with different BK constructs

Peak current resulting from activation of an inactivating channel is expected to be less than the maximum current that would arise from all channels being open simultaneously. The reduction in peak current can arise from a number of factors. First, channels may already occupy inactivated states prior to the depolarizing voltage step. Second, inactivation of some channels may occur during the rising phase of current activation prior to the time of peak current. In this case, the extent to which peak current may underestimate the true maximum current differs dependent on whether the inactivation process may be dependent on opening first occurring (coupled model) or that inactivation can occur without channel opening being driven only by depolarization (uncoupled model). For the simplest formulation of coupled and uncoupled models, to evaluate the expected reduction of peak current relative to the measured maximum activatable BK current in a patch only requires measurement of the inactivation time constant and then, following removal of inactivation by trypsin, measurement of the activation time constant and the maximum current amplitude during saturating activation for the population of channels in the patch.

Although the extent to which BK inactivation is coupled or independent of activation has not been addressed unlike the situation for Shaker (Zagotta et al., 1989), here we assume two limiting analytic approximations, beginning with the assumptions that for BK currents the activation time course can be reasonably well-described by a single exponential function (Horrigan and Aldrich, 2002) and that the stimulation conditions are such that return from O states to C states is minimal. For the first case, we assume that inactivation is uncoupled from activation, i.e., it occurs independent of activation, being driven solely by depolarization. This defines a potential lower limit on peak current amplitude that might result from inactivation occurring during the rising phase, and is described by the following sort of Hodgkin-Huxley activation-inactivation formalism, 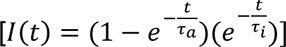. with τ_a_ and τ_i_ reflecting activation and inactivation time constants, respectively. For the second approximation, we assume that inactivation can only occur following channel opening, i.e., inactivation is fully coupled to opening, as defined by a linear scheme of C→O→I. In this case, the explicit analytic equation is: 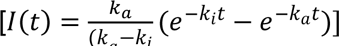, with *k*_a_ and *k*_i_ being activation and inactivation rates, respectively. In either case, whether passage through closed states is approximated by a single exponential time course or might involve multiple transitions, as long as the function reasonably approximates the current rise time, the expected consequences of the two inactivation formalisms are reasonably described.

### Simulations of two-step inactivation behavior

Simulations utilized IonChannelLab software (v. 1.0.5.3) (Santiago-Castillo et al., 2010). Three primary stimulus protocols were utilized, one to mimic that used in Fig. 3A for current activation and basic tail currents, a second to mimic that used in Fig. 7 to look at voltage-dependence of tail current following inactivation, and then a third identical to the steady-state inactivation protocol used for Fig. 4D. A population of 1000 channels was assumed and currents were simulated at a sampling interval of either 20 or 100 μs per point. We used a simplified channel activation model as given in Scheme 1. The basic strategy was to determine whether, within the context of a simplified model, could changes in a minimal subset of parameters account for the experimental observations.

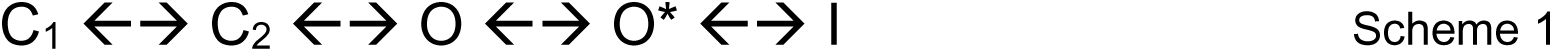

The crux of the 2-step inactivation model is that the transition from O to O* is the rate limiting step in the onset of inactivation, while the O*-I equilibrium involves transitions much faster than can be measured with available techniques. BK channel activation was approximated with three states, C_1_ ←→ C_2_ ←→ O, with rates and voltage-dependencies chosen to generate GVs and activation/deactivation kinetics that mimic those for the four BK constructs (WT, RKK3Q, RKK3E, and RKK3V) when in association with D20Am after removal of inactivation by trypsin (D20Am-IR). Based on the experimentally measured onset of inactivation at positive potentials (∼16 ms), the O→O* transition was set at 60/s unless otherwise indicated. The rates that underly the O*←→I process were taken from previous work based on a β-analysis of the reduced current level openings produced by the β3a(1-20) peptide on WT BK channels (Gonzalez-Perez et al., 2012) The voltage-dependence of the O*←→ I equilibrium was also confirmed to be similar in Fig. 6D-E for single channels arising from D20Am expressed with RKK3V. Negative to −100 mV, D20Am+RKK3V channels begin to exhibit marked inward current rectification negative resulting in a rollover in conductance values in contrast to those for the β3a(1-2) peptide. This rollover arises from the marked inward current rectification at negative voltages for BK channels expressed with β2 or β3 subunits that arises from the extracellular domain of the β subunits (Zeng et al., 2003).

## Results

### A potential gate in the side portal region of BK channel

Previous work suggests that BK β-subunit cytosolic N-terminal inactivation segments likely reach their inactivation sites by passage through a side portal formed between the four linkers connecting the transmembrane domain and the CTD of a BK channel (Zhang et al., 2006; Zhang et al., 2009). To evaluate elements that might define possible pathways, we attempted to define aqueous volumes in closed and open BK channel structures that might permit peptide chains to access the central cavity. Based on the globular dimensions of the N-terminal hydrophobic motif (MFIW) of the BK β2 subunit (Bentrop et al., 2001) that is required for β2-mediated inactivation (Xia et al., 2003), we computed the aqueous volume accessible to a particle 8 Å in diameter in two available hSlo1 Cryo-EM structures (Tao and MacKinnon, 2019) using program PrinCCes (Czirjak, 2015). In the Ca^2+^-bound structure (Fig. 1A), an aqueous accessible volume can be seen to extend laterally from the central cavity to the outside of the PGD. In contrast, in the apo structure clear gaps separate the aqueous volume within the central cavity of the PGD from the aqueous volumes outside the PGD (Fig. 1B). Closer inspection shows that the gap in the apo structure results from intrusion of the side chains of three consecutive basic residues (R_329_K_330_K_331_; RKK) in the N-terminus of BK C-linker (Fig. 1C). This suggests that, in the closed channel, RKK may hinder access of β inactivation segments to the BK central cavity. The potential importance of the RKK residues in influencing state-dependent access of β subunit N-termini to the central cavity can also be inferred from examination of recent structures of a β2NT-β4 chimera in association with hSlo1 (Agarwal et al., 2025).

**Fig. 1.**
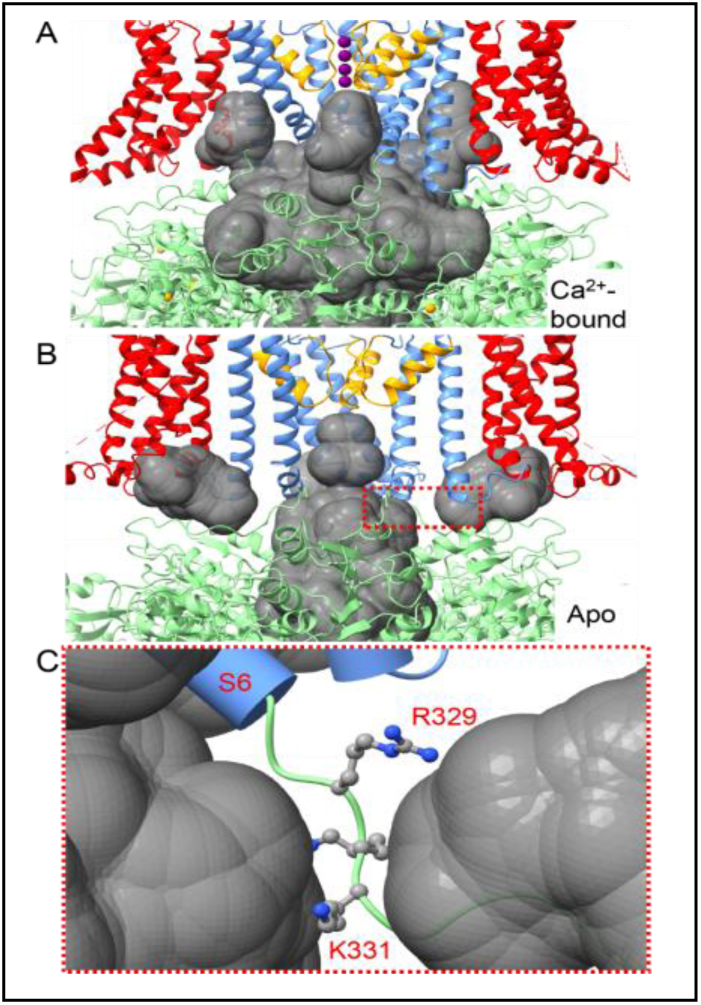
BK side portal visualization. (A) The Ca^2+^-bound hSlo1 structure (PDB: 6V3G) and the interior aqueous space (gray) accessible to a particle of 8-Å in diameter. The hSlo1 structure is rendered as ribbon, with VSD, PGD, and CTD colored in red, blue, and green, respectively. The selectivity filter is colored in orange with K^+^ rendered as purple spheres. The orange spheres in the CTD are Ca^2+^ ions. (B) The apo hSlo1 structure (PDB: 6V38) and the aqueous space (gray) accessible to a particle of 8-Å in diameter. The color scheme is identical to that used in panel A. The red box highlights the gap between the central volume and the volume outside the PGD. (C) A blowup view of the region highlighted by red box in panel B, with the C-linker basic triplet (RKK) rendered as ball-and-stick and S6 as cylinder.

### Framework for evaluation of the potential role of the RKK triplet in perturbing inactivation by BK β3 and β2 subunits

We utilized three mSlo1 mutants: RKK3Q, RKK3E, and RKK3V (Fig. 2A). To probe inactivation, we coexpressed each Slo1 construct with the D20Am β3a:β2 chimeric construct (β3a(1-34):β2(2-235)), in which the mouse β3a (mβ3a) N-terminus (Fig. 2B) replaces the N-terminus in a human β2 (hβ2) subunit (Gonzalez-Perez et al., 2012). For this work, the D20Am construct offers several advantages. It retains all the relevant functional characteristics of β3a-mediated inactivation, including the prominent features of two-step inactivation (Scheme 1 as in Fig. 2C). The reason for using the D20Am chimera over a native β3a construct is that it exhibits reduced current rectification properties known to arise from β3 subunit extracellular loops (Zeng et al., 2003). D20Am produces inactivation-dependent slow tail current (Fig. 2D) that is removed following trypsin removal of inactivation (Fig. 2E). Furthermore, single BK+D20Am channels exhibit prolonged openings of reduced average current amplitude (compared to full openings) that occur immediately upon repolarization (Fig. 2F). Trypsin removal of inactivation also abolishes the slow tail openings (Fig. 2F). The tail openings of reduced amplitude exhibit the following features. Opening is essentially instantaneous following repolarization. The reduction of current amplitude relative to a normal BK tail opening is relieved as the membrane voltage becomes more negative (Fig. 2G), approaching the amplitude of normal tail openings at very negative voltages. The largely complete inactivation at positive voltages and essentially instantaneous opening upon repolarization to a reduced conductance level can be explained by rapid transitions between two states, a preinactivated open state (O*) and a very brief non-conducting state (I) in accordance with the two-step inactivation scheme (Fig. 2C) (Zeng et al., 2007; Gonzalez-Perez et al., 2012). The voltage-dependence of the O*←→I equilibrium is such that occupancy in I is strongly favored with depolarization, but O* is favored with hyperpolarization. The idea that D20Am tail currents largely reflect a population of channels defined by the voltage-dependence of the O*/I equilibrium underlies some of results and analysis presented below, and new results further bolster the idea that the general framework of the 2-step model is a reasonable explanation for D20Am-mediated inactivation. Guided by these considerations, we can now present results with mutated RKK constructs.

**Figure 2.**
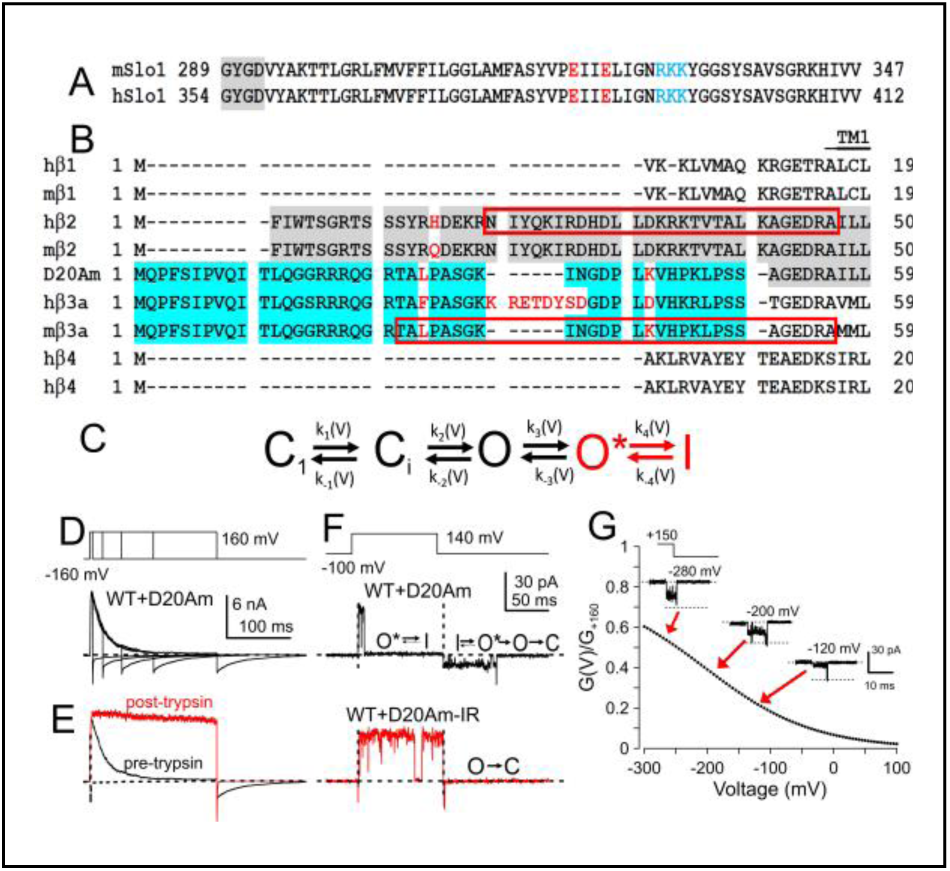
Constructs and β3a behavior. **(A)** Amino acid sequences of mouse and human Slo1 spanning the GYGD selectivity filter, the S6 segment, and the so-called C-linker to the beginning of RCK1. Positions of the RKK triplet and E321/E324 (mouse) are highlighted. **(B)** Sequences of human and mouse β subunit N-termini. **(C)** Two-step inactivation scheme. **(D)** WT+D20Am currents were activated in 10 μM Ca^2+^ by steps to +160 mV of varying duration. As inactivation develops, the tail currents become markedly prolonged, greatly exceeding the net current flux expected from inactivated channels simply returning to an open state before closing. **(E)** Trypsin removes inactivation and abolishes the slow tail current. **(F)** Activation of a single inactivating BK+D20Am channel by the indicated voltage protocol with 10 μM Ca^2+^ before (trace in black) and after (trace in red) application of trypsin to remove inactivation. Note the tail opening following inactivation occurs immediately upon repolarization and opens to a reduced current level. Following removal of inactivation by trypsin (WT+D20Am-IR), only brief, large amplitude tail openings are observed. **(G)** Examples of O*-I tail openings at different voltages produced by β3a(1-20) L-peptide, along with curve reflecting voltage-dependence of O*/(O*+I) relationship based on β3a peptide action (from (Gonzalez-Perez et al., 2012)).

### RKK mutations markedly reduce inactivating outward current and generate a largely persistent inward current

Each of the three RKK3 mutant construct we have examined forms functional BK currents, with GV relationships at 0 Ca^2+^ (Table S1) very comparable to those reported previously (Tian et al., 2019). To assess the gating range of the WT and RKK mutant channels when coexpressed with D20Am (Fig. S1), we first measured GV curves for each construct after trypsin removal of inactivation (Fig. 3A2,B2,C2,D2 for WT, RKK3Q, RKK3E, and RKK3V, respectively). This provided information regarding the range of voltages at a given [Ca^2+^] over which each construct in association with D20Am would be expected to occupy mainly closed states both during prepulses to −160 mV and following repolarizations to −160 mV. The GVs defined the range of voltages over which saturating macroscopic current activation occurs at a given Ca^2+^. Based on these GVs, we selected 10 μM Ca^2+^ for most protocols for WT, RKK3Q, and RKK3Ewhen expressed with D20Am, since at −160 mV there is little activation. For RKK3V+D20Am, the activation is more negatively shifted (Fig. 3D), such that we have generally used 0 Ca^2+^ so that prepulses to −160 mV minimize channel openings. These choices were confirmed in single channel patches in which 10 μM Ca^2+^ rarely opens RRK3Q+D20Am channels at −160 mV after removal of inactivation (Fig. S1E), while with 0 Ca^2+^ RKK3V+D20Am channels rarely open at −160 mV (Fig. S1F).

**Figure 3.**
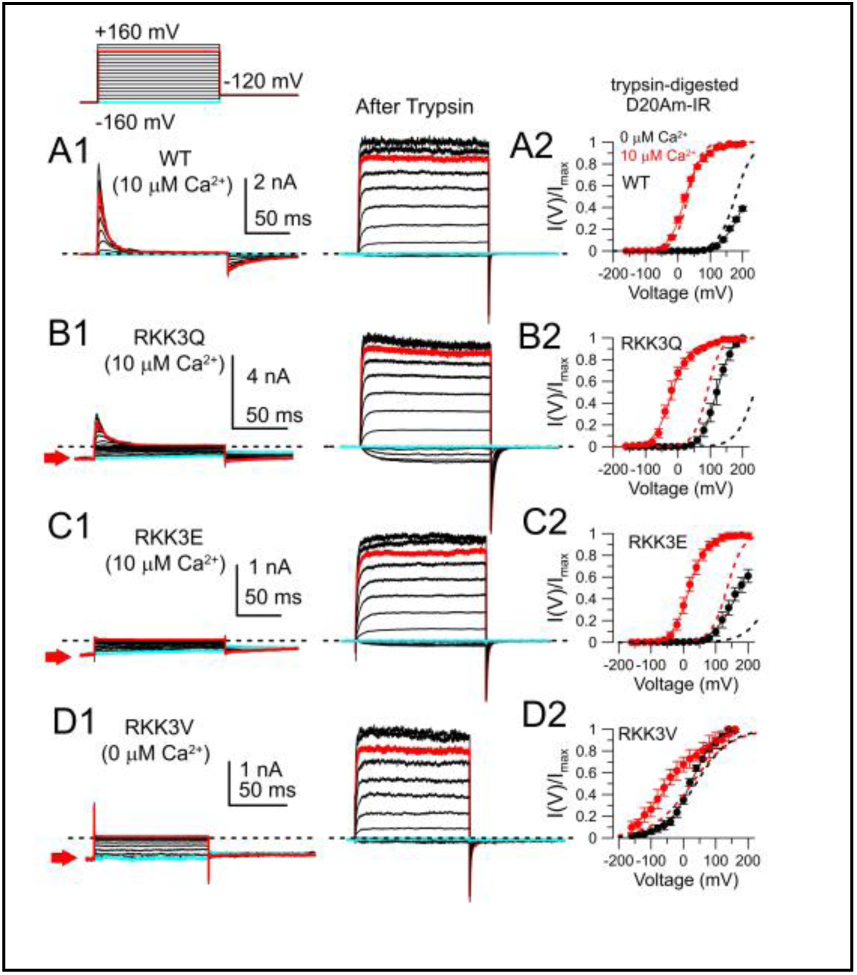
RKK mutations increase occupancy of inactivated states at negative potentials. (A1) A standard activation protocol (inset) elicited current activation and inactivation of BK+D20Am channels before (left) and after (right) trypsin digestion. Trypsin abolishes the slow tail following repolarization from inactivation. Red trace: +120 mV; cyan trace: −160 mV. 10 μM Ca^2+^. (A2) GV curves from WT+D20Am-IR channels. Dotted lines correspond to GVs for WT currents without coexpression with D20Am (values in Table S1). The Boltzmann fits to the D20Am-IR curves are : V_50_ = 211.1 ± 5.1 mV, *z* = 0.71 ± 0.09*e* (0 Ca^2+^); V_50_ = 22.1 ± 2.1 mV, z = 1.01 ± 0.07*e* (10 μM Ca^2+^). (B1) Currents for RKK3Q+D20Am show reduced inactivating outward current, but a persistent inward current at negative potentials (highlighted by red arrow on left), which disappears after trypsin digestion. 10 μM Ca^2+^. (B2) GV curves for RKK3Q+D20Am-IR channels, with dotted lines for RKK3Q alone: V_50_ = 120.7 ± 4.1 mV, *z* = 1.04 ± 0.13*e* (0 Ca^2+^); V_50_ = −20.2 ± 3.1 mV, *z* = 0.96 ± 0.10*e* (10 μM Ca^2+^). (C1) For RKK3E+D20Am currents, time-dependent inactivation and opening at depolarized potentials is completely absent, despite persistent inward current at negative potentials. 10 μM Ca^2+^. (C2) GV curves for RKK3E+D20Am-IR: V_50_ = 172.2 ± 3.4 mV, *z* = 0.65 ± 0.06*e* (0 Ca^2+^); V_50_ = 17.1 ± 2.4 mV, *z* = 1.00 ± 0.08*e* (10 mM Ca^2+^). (D1) For RKK3V+D20Am, even with 0 Ca^2+^, there is no outward current activation or inactivation, although persistent inward current is even larger as a percentage of maximum available BK current. (D2) GV curves for RKK3V+D20Am-IR: V_50_ = 23.8 ± 6.2 mV, *z* = 0.64 ± 0.10*e* (0 Ca^2+^); V_50_ = −47.0 ± 8.5 mV, *z* = 0.44 ± 0.06*e* (10 μM Ca^2+^).

Macroscopic currents from WT (Fig. 3A) and RKK-mutated (Fig. 3B-D) BK α subunits coexpressed with D20Am were evoked from inside-out patches by test pulses from −160 to +160 mV following a 200 ms pre-pulse to −160 mV. After recording the inactivating currents (Fig. 3A-D, left panels), each patch was treated with trypsin to remove inactivation (Fig. 3A-D, right panels). This allows definition of the maximum BK conductance in a patch. We note that, for the RKK mutant constructs, the peak current estimated by trypsin digestion is likely somewhat underestimated, given that there is some residual fast flickery gating following trypsin digestion as shown below in single channel records (e.g., Fig. S1E,F). This likely reflects fast block by residual trypsin-digested N-termini that produces a small decrease in apparent maximum conductance.

For WT+D20Am BK current recorded in 10 μM [Ca^2+^]_in_, the outward inactivating current evoked by the step to +160 mV peaks at around 85% of the maximal conductance (defined following removal of inactivation by trypsin) before decaying close to the zero current level (Fig. 3A), indicating that most WT channels open before they inactivate. Repolarization during inactivation results immediately in the signature slow tail current involving the β3a N-terminus. After digestion by trypsin, the slow tail is completely eliminated together with inactivation. In contrast, currents arising from RKK3Q coexpression with D20Am exhibit two noticeable differences from the WT inactivating BK current (Fig. 3B). First, the peak outward current of RKK3Q is reduced to about 35% of the maximum current after trypsin digestion. Second, with the 200 ms prepulses to −160 mV, there is an appreciable amount of inward current at potentials negative to −100 mV (Fig. 3B, cyan trace: −160 mV). Qualitatively, the reduction of peak outward current might be interpreted to indicate that at least twice as many RKK3Q channels are inactivated at the peak of the outward current in comparison to WT. However, this reduction might occur either from inactivation during the rising phase of current or from inactivation occurring prior to the depolarization. Macroscopic currents for the other two RKK mutants are shown in Fig. 3C (RKK3E, 10 μM [Ca^2+^]_in_) and Fig. 3D (RKK3V, 0 μM [Ca^2+^]_in_). Similar to RKK3Q, both RKK3E and RKK3V when coexpressed with D20Am result in sustained inward current at negative potentials. This inward current is removed by trypsin treatment (Fig. 3C-D). However, unlike either RKK3Q or WT channels, there is almost no outward current in RKK3E or RKK3V channels expressed with D20Am until after trypsin digestion. Yet, this pronounced suppression of voltage-activated outward current is associated with a measurable inward conductance at −160 mV and other negative potentials.

Simplistically, the reduction in depolarization-activated peak outward current in the RKK3X+D20Am constructs might be interpreted to suggest that there is an increased likelihood that channels already reside in inactivated states at the end of the 200 ms prepulse at −160 mV preceding depolarization. Thus, one interpretation of the reduction of outward current would be that the RKK mutations are favoring inactivation directly from closed states. However, the inward current at −160 mV observed for each of the RKK mutant constructs complicates interpretation of the state-occupancies of channels during and at the end of the −160 mV prepulse, i.e., are channels occupying closed states? Below we attempt to systematically address this question.

For the RKK3Q, RKK3E, and RKK3V mutants when expressed with D20Am, the tail currents arising from repolarization to −160 mV all appear to be more prolonged versions of the WT+D20Am tail current. Based on this idea, in accordance with the 2-step inactivation scheme, we suggest that the inward current reflects some fraction of channels undergoing rapid oscillations between O* and I states (O*←→I). If so, depolarization from −160 mV would immediately result in a new O*/I equilibrium favoring occupancy predominantly in I. Thus, any channels occupying the O*-I ensemble at negative potentials will contribute not only an inward current at negative potentials, but also will not be available to contribute to channel activation and inactivation upon depolarization. Yet, the question remains whether any of the reduction in outward current arises from channels that directly inactivate from closed states, rather than channels in the ensemble of O*-I states. Much of the subsequent experiments are designed to evaluate whether the RKK mutations are permitting direct inactivation from closed states.

### Changes in activation or inactivation rates in RKK/3X+D20Am channels do not account for reduction in peak outward current

Differential changes in the rates of inactivation onset and activation in the RKK-mutated channels might contribute to a fraction of RKK-mutated channels entering inactivated states following depolarization, but preceding current peak. To address this possibility, following removal of inactivation by trypsin, we measured activation time constants as a function of membrane potential over the range of +60 to +160 mV (Fig. 4A. left). Similarly, we measured inactivation time constants at +160 mV prior to trypsin digestion (Fig. 4A, right). For RKK3E, only a very small transient current was observed; inactivating currents could not be meaningfully measured for RKK3V.

**Figure 4.**
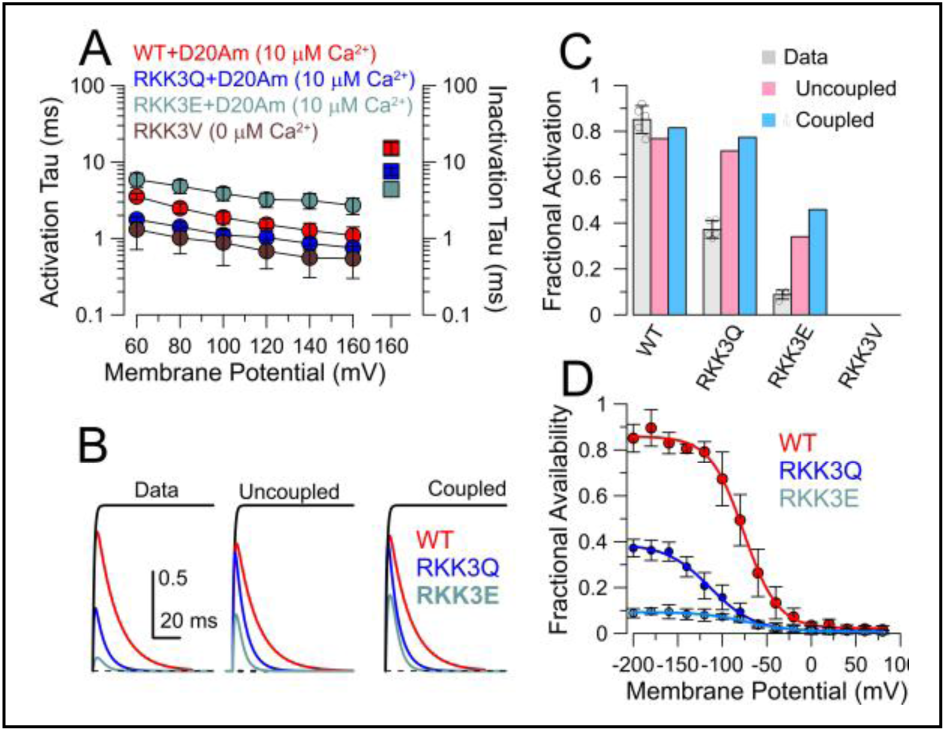
RKK mutations increase occupancy of inactivated states at negative potentials. **(A)**. Left axis, activation time constants for each construct following trypsin removal of inactivation are plotted as a function of command potential. Right axis, inactivation time constants for three of the constructs measured at +160 mV are shown on the right. **(B)** Measured (Data) and predicted (Uncoupled and Coupled) current activation and inactivation time courses normalized to maximum current after trypsin digestion are displayed for WT, RKK3Q, and RKK3E, based on the measured activation and inactivation time constants from panel A. **(C)** Measured (gray) and predicted fractional activation for each construct following a prepulse to −200 mV. **(D)** A 300 ms prepulse to voltages from −200 to +80 mV, followed by current activation at +160 mV, was used to generate a steady-state inactivation curve for the indicated constructs. Peak current at +160 mV measured following each prepulse voltage was normalized to peak current at +160 mV following removal of inactivation. Solid lines are fits of a Boltzmann function. Fit parameters were: for WT (n=5 patches), h_∞_=0.86±0.02; V_50_ = −74.8±2.1 mV, *z*=1.2±0.1e; for RKK3Q (n=5), h_∞_=0.40±0.02; V_50_=-113.1±3.8 mV; *z*=0.94±0.01e; for RKK3E (n=4), h_∞_=0.10±0.01; V_50_=-68.2±7.2 mV; z=0.75±0.12e. Comparisons of fractional availability at −200 mV among each construct with ANOVA with Tukey’s multiple comparisons yielded, for WT vs. RKK3Q, P<0.0001; for WT vs. RKK3E, P<0.0001; for RKK3Q vs RKK3E, P=0.0002.

The extent to which channels may inactivate during the rising phase of current activation is influenced by the specific relationship between activation and inactivation. For BK, the extent to which inactivation is coupled or independent of activation has not been addressed unlike the situation for *Shaker* (Zagotta et al., 1989). Therefore, for this evaluation, here we assume two limiting analytic approximations (see Methods), beginning with the assumption that, for BK currents, the activation time course can be reasonably well-described by a single exponential function (Horrigan and Aldrich, 2002). We also assume that, at +160 mV, once channels enter the O*-I ensemble they do not return to O therefore allowing it to be approximated by O→I. For the first case, we assume that inactivation is “uncoupled” from activation, i.e., inactivation can be initiated by depolarization even without channel opening. This is described by the following Hodgkin-Huxley activation-inactivation formalism, 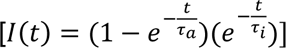. For an inactivation process uncoupled from activation, the activation and inactivation time constants lead to estimates of 0.77, 0.71, and 0.34 for the predicted fraction of peak current expected for WT, RKK3Q, and RKK3E (Fig. 4B, middle panel, 4C). For an inactivation process in which inactivation depends on channels first opening (”coupled”), based on a linear scheme of C→O→I, when return from O to C is minimal as might be expected with strong depolarizations, the explicit analytic equation is: 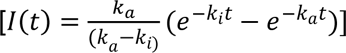. Based on the measured time constants, this predicts peak current (Fig. 4B) with amplitudes of 0.82, 0.77, and 0.45, for WT, RKK3Q, and RKK3E, respectively). For WT currents, both models make predictions similar to what is observed experimentally, suggesting that there may be little inactivation of WT channels at −160 mV prior to the depolarization (Fig. 4B,C). However, for RRK3Q and RKK3E, the experimentally observed reduction in peak current is significantly more than the predictions from either the coupled or uncoupled model of inactivation. This suggests that approximately 40% and 60% of RKK3Q and RKK3E channels, respectively, may already reside in inactivated states prior to depolarizing pulses. The absence of appreciable outward current for RKK3V+D20Am precludes this kind of calculation, but depolarization from −160 mV for RKK3V+D20Am channels rarely activates normal channel opening. These considerations again suggest that the RKK triplet mutations examined here increase the likelihood that channels at the end of the prepulse to −160 mV reside in states that are unable to transition to normal openings. By standard convention, that would mean the channels are in inactivated states. Whether that solely reflects occupancy in the O*-I ensemble or some other inactivated state is addressed below.

### At negative potentials, RKK mutant channels occupy closed states that are in a voltage-independent equilibrium with an inactivated state

Since our patch recordings typically use a holding potential of 0 mV before initiation of protocols, 200 ms at −160 mV may be insufficient to return channels to resting states. Therefore, we generated steady-state inactivation curves for each construct over a range of voltages down to −200 mV, while increasing the prepulse to 300 ms (Fig. 4D). For each patch we normalized the peak of inactivating currents to the maximum available conductance defined after trypsin. Thus, the steady-state inactivation curves potentially define the true fractional availability of a given channel to be activated, with some reduction in the peak current expected from the fraction of channels that inactivate following depolarization, but preceding current peak. For all three constructs, each exhibited a specific limiting fractional availability (ℎ_∞_) that was unchanged between −160 and −200 mV at values comparable to the peak fractional activation described above (Fig. 4B-C). The V_50_ of steady-state inactivation was left-shifted for RKK3Q relative to WT and RKK3E. For RKK3V, the longer conditioning step to −200 mV failed to unmask any activation of outward current. More negative voltages of longer duration were not well-tolerated by the patches. A concern with these considerations is that perhaps the prepulse duration is insufficient to define true fractional availability. Experiments below suggest that, at −160 and −200 mV, persistent current is near steady-state within about 200 ms.

That RKK3Q, RKK3E, and RRK3V each reach a limiting, but differing, asymptote suggests that each channel reaches a different voltage-independent fraction of channels that are available to open normally and inactivate following depolarization. Some of the reduction in fractional availability is expected to reflect channels in the O*-I ensemble at −160 mV, but we suggest that, for RKK3Q and RKK3E, there may also be a fraction of channels that are inactivated from fully resting closed states. These ideas are developed further below.

Typically, steady-state inactivation curves are used to define a voltage of half availability (V_50_) and are simply normalized to the peak inactivating current measured during a given set of depolarizations. Here we have benefit of an independent measure of the total current expected for the population of channels in a patch when inactivation is removed, something that cannot readily be achieved for most other inactivating channels (although see (Ayer and Sigworth, 1997)). The differences in limiting asymptote therefore provide a direct and essentially model-independent measure of differences in voltage-independent fractional occupancy of channels in activatable versus inactivated states.

### Single channel behavior of D20Am+RKK mutant constructs supports the idea that persistent current reflects channels in O*-I states

To probe the single channel underpinnings of the persistent inward current, we examined single channel patches with a stimulation protocol similar to that used in Figure 3. We designate the initial prepulse to −160 mV as segment S1, the step to +140 mV as S2, and the repolarization to −160 mV as S3 (top of Fig. 5). For WT+D20Am channels (Fig. 5A), depolarization to +140 mV resulted in openings that then inactivated in 608 of 793 trials and 37 openings that remained open for the duration of the depolarization (n= 5 patches) (Table 1). Repolarization of channels that were inactivated at end of S2 usually resulted in immediate channel opening to a current level intermediate between fully open and fully closed as described previously for the O*-I behavior (Zeng et al., 2007; Gonzalez-Perez et al., 2012). Following trypsin removal of inactivation, WT+D20Am-IR channels remained open for the duration of the depolarization; after repolarization, brief tail openings of larger amplitude could occasionally be observed, although at −160 mV most tail openings are too brief to be resolved (e.g. Fig. 5E). Ensemble averages of the single channel events both before and after trypsin (Fig. 5A, bottom) mirrored the macroscopic behaviors shown above. For RKK3Q+D20Am (Fig. 5B), depolarization to +140 mV occasionally elicits normal openings followed by inactivation, but at lower likelihood than for WT+D20Am (125 openings in 359 sweeps from 6 patches) (Table 1). However, in contrast to WT+D20Am, the RKK3Q+D20Am tail current openings that occurred immediately upon repolarization were more prolonged, often persisting for the duration of the repolarizing step. Trypsin also removed the prolonged persistent openings, and ensemble averages mirrored the macroscopic currents (Fig. 5B). For RKK3E+D20Am (Fig. 5C), very few sweeps revealed channel openings during the depolarization to +140 mV (10 openings in 330 sweeps from 8 patches (Table 1)), while channel openings of reduced amplitude were also immediately observed upon repolarization. Finally, for RKK3V+D30Am (Fig. 5D), single channel openings during steps to +140 mV were also rarely seen (12 of 588 sweeps from 8 patches (Table 1)), while repolarization was invariably associated with immediate reopening of reduced amplitude level that persisted for the duration of the repolarization. A characteristic of the single channel openings for RKK3Q, RKK3E, and RKK3V following trypsin digestion of D20Am was the flickery nature of the openings (e.g., Fig. S1E-F). As noted above, we suggest that residual digested N-termini are able to mediate a rapid flickery block behavior that does not occur with the WT RKK sequence, perhaps reflecting an interaction of the residual N-terminus with something affected by mutation of RKK residues.

**Figure 5.**
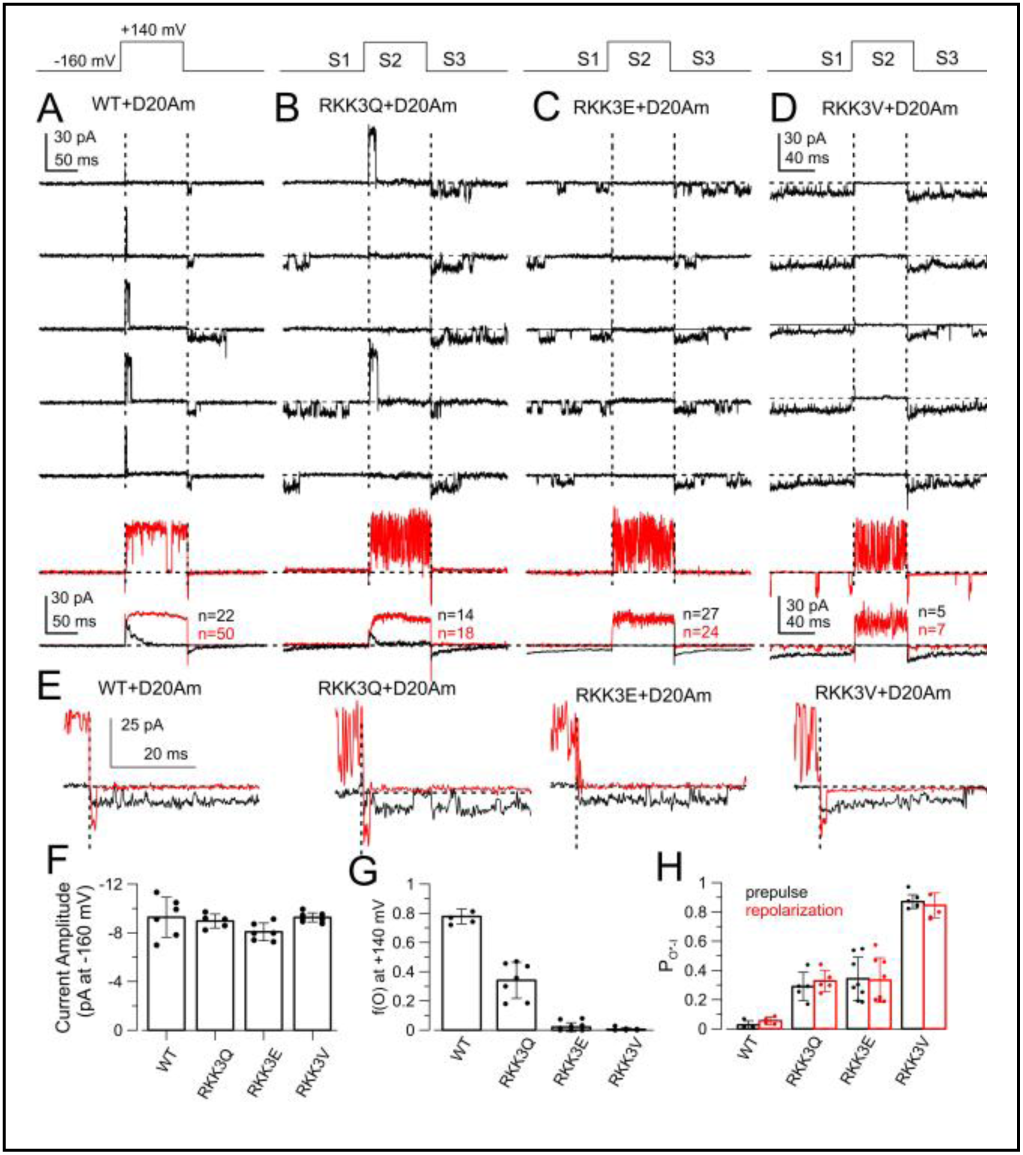
D20Am-RKK mutant channels exhibit increased single channel activity at negative potentials, with reduced likelihood of opening during depolarizing steps. **(A)** A WT Slo1+D20Am single channel was activated with the indicated voltage protocol and 10 μM Ca^2+^. Across top, Hyperpolarizing prepulse, depolarization, and repolarization in stimulation protocol are designated S1, S2, S3, respectively. Most trials result in channel opening during depolarization following by inactivation. Repolarization results immediately in long duration tail current openings. Red trace in 5^th^ row from top shows activation after trypsin removal of inactivation highlighting the briefer and larger amplitude tail opening after removal of inactivation. On the bottom, ensemble averages for traces before (black) and after (red) removal of inactivation are plotted with the number of sweeps in the average on the panel. (B) D20Am+RKK3Q channels exhibit frequent low conductance openings at −160 mV, while depolarization results in less likely normal activation and inactivation. Prolonged low conductance openings for D20Am+RKK3Q channels disappear following trypsin, while the single channel amplitude returns to WT levels (red traces). (C) For D20Am+RKK3E, depolarization rarely produces normal openings and inactivation, while reduced conductance openings are observed at −160 mV. Trypsin abolishes the persistent inward openings and abolishes the slow tail current. (D) RKK3V+D20Am channels rarely produce normal openings and inactivation with depolarization, but generate persistent inward openings that are quickly abolished after D20Am N-terminal digestion by trypsin. (E) Higher resolution examples of tail current low amplitude openings showing the similarity in amplitude for each of the four constructs. (Red) Tail openings after removal of inactivation. RKK3E tail openings after digestion were too brief to be resolved. (F) Mean (±SD) and individual estimates of amplitudes of open current level following repolarization at −160 mV (WT, N=6; RKK3Q, N=5; RKK3E, N=6; RKK3V, N=8). For all comparisons of amplitudes among constructs, P>0.1194 for ANOVA with Tukey’s correction for multiple comparisons. (G) Likelihood of depolarization to 140 mV producing an opening following 150 ms at −160 mV is plotted for 5 WT+D20Am patches (N= 793 trials), 6 RKK3Q+D20Am patches (N=359 trials), 8 RKK3E+D20Am patches (N=330 trials), and 8 RKK3V+D20Am patches (N=588 trials). (H) Probability of each type of channel occupying low conductance channel openings (designated P_O*-I_) is displayed, for both before (black) and after (red) the depolarization. These estimates are not intended to reflect true steady state P_O*-I_ at −160 mV.

**Table 1.** Single Channel Behavior Immediately Preceding (S1) and After (S2) Depolarization.

|  | **Evet Occurrences** | **WT (n=5)**<br>**150 ms pp** | **WT (n=6)**<br>**250 ms pp** | **RKK3Q (n=6)** | **RKK3E**<br>**(n=8)** | **RKK3V**<br>**(n=8)** |
| --- | --- | --- | --- | --- | --- | --- |
| 1 | N | 793 | 396 | 359 | 330 | 588 |
| 2 | C<sub>S1</sub> | 790 | 396 | 299 | 256 | 119 |
| 3 | O*<sub>S1</sub> | 3 | 0 | 59 | 74 | 469 |
| 4 | O<sub>S1</sub> | 0 | 0 | 1 | 0 | 0 |
| 5 | C<sub>S2</sub> (never opens) | 148 | 41 | 234 | 320 | 576 |
| 6 | O<sup>I</sup><sub>S2</sub> inactivation | 608 | 320 | 125 | 10 | 12 |
| 7 | O<sup>S</sup><sub>S2</sub> sustained | 37 | 35 | 0 | 0 | 0 |
| **Association of S2 events with channel status at end of S1** |  |  |  |  |  |  |
| 8 | O*<sub>S1</sub>→C<sub>S2</sub> | 3 (0.003) | 0 (0.00) | 59 (0.164) | 74 (0.224) | 469 (0.582) |
| 9 | O*<sub>S1</sub>→O<sub>S2</sub> | 0 (0.00) | 0 (0.00) | 1 (0.003) | 0 (0.00) | 0 (0.00) |
| 10 | C<sub>S1</sub>→C<sub>S2</sub> | 145 (0.183) | 41 (0.103) | 175 (0.488) | 246 (0.745) | 107 (0.375) |
| 11 | C<sub>S1</sub>→O<sub>S2</sub> | 645 (0.813) | 355(0.896) | 124 (0.345) | 10 (0.030) | 12 (0.042) |
| **Fraction of trials with S2 opening ((r6+r7)/r1)** |  |  |  |  |  |  |
| 12 | O<sub>S2</sub>/N | 0.813 | 0.896 | 0.348 | 0.03 | 0.020 |
| **Fraction that fail to open in S2 when S1 terminates in O\* (r8/(r8+r9))** |  |  |  |  |  |  |
| 13 | (O*<sub>S1</sub>)→C<sub>S2</sub>/ O*<sub>S1</sub> | 1.0 (3/3) | N.A. | 0.983 (59/60) | 1.0 (74/74) | 1.0 (469/469) |
| **Fraction of sweeps in inactivated states at end of S1 (1-r12)** |  |  |  |  |  |  |
| 14 | O*<sub>S1</sub> + (C<sub>S1</sub>→C<sub>S2</sub>) /N | 0.187 | 0.103 | 0.652 | 0.970 | 0.980 |
| **Fraction of long closures at end of S1 that likely represent inactivated states (r10/r2)** |  |  |  |  |  |  |
| 15 | (C<sub>S1</sub>→C<sub>S2</sub>)/C<sub>S1</sub> | 0.184<br>(145/790) | 0.103<br>(41/396) | 0.585<br>(175/299) | 0.961<br>(246/256) | 0.899<br>(107/119) |

Nomenclature: S1 and S2 denote segments S1 and S2 in activation protocol
Formalism "O\*<sub>S1</sub>→C<sub>S2</sub>" indicates C<sub>S2</sub> events that occur subsequent to O\*<sub>S1</sub> event at end of S1.
C<sub>S1</sub>: Channel is closed at end of S1
O\*<sub>S1</sub>: Channel is in a reduced conductance burst at end of S1
C<sub>S2</sub>: channel never opens during S2
O<sub>S2</sub>: channel opens during S2
Row 3. Likelihood that S1 ends in O\* increases from WT<3Q<3E<3V. O\*<sub>S1</sub> events vs. non-O\*<sub>S1</sub> events. Fishers exact with Bonferroni. WT1 vs. WT2, P=0.555; WT vs. 3Q, P<0.0007; WT vs 3E, P<0.0007; WT vs. 3V, P<0.0007; 3Q vs. 3E, P=0.0532; 3Q vs. 3V, P<0.0007; 3E vs. 3V, P<0.0007.
Row 13. O\*<sub>S1</sub> invariably results in C<sub>S2</sub> for all constructs.
Row 14. Occupancy of an inactivated state (O\*<sub>S1</sub> or C<sub>S1</sub>→C<sub>S2</sub>) at end of S1 increases in order of WT<3Q<3E~3V. WT1 vs. WT2, P=0.0014; WT1 vs. 3Q, P<0.0007; WT1 vs. 3E, P<0.0007; WT1 vs. 3V, P<0.0007; 3Q vs. 3E, P<0.0007; 3Q vs 3V, P<0.0007; 3E vs. 3V, P=0.2156.

Examination of WT+D20Am tail openings in the S3 segment (Fig. 5E, left) both before and after trypsin emphasize the nature of the reduced current level during the O*-I burst relative to a normal tail opening. The openings for RKK3Q+D20Am (Fig. 5E, middle left, Fig. 5F) suggest the amplitudes of the presumed O*-I bursts are similar to those for WT, while the durations of the bursts are longer (Fig. 5B). The single channel current level for the RKK3E+D20Am channels was also indistinguishable from WT (Fig. 5E, middle right, Fig. 5F); given the brevity of the RKK3E+D20AM-IR tail openings after trypsin, we were unable to identify a fully resolved tail opening in the set of sweeps we recorded. Finally, RKK3V+D20Am single channel openings following repolarization were also of similar amplitude to those of the other RKK constructs (Fig. 5E, right, 5F). Based on the similarity of amplitudes of tail current opening for each of the four complexes (Fig. 5F), we suggest that the openings likely reflect the same underlying behavior (reflecting O*-I bursts) defined by interaction of the β3a N-terminus within the BK central cavity. The different durations of the tail openings and the presence of persistent current would seem likely to arise from differences in the affinity of interaction of the β3a N-terminus within the central cavity, influenced by the specific triplet of residues.

### Failure to open during S2 depolarizations can follow either from occupancy of channels in the O*-I ensemble or from a longer lived non-conducting state

The single channel records permit an evaluation of the likelihood of observing a given behavior after a voltage-step contingent upon channel status just preceding the voltage step. Fig. 6A illustrates the various types of channel behaviors that can be seen at the end of segment S1 in association with channel status at the beginning of segment S2. At the end of S1, a channel may be either closed (C_S1_), open (O_S1_), or in an O*-I burst (O*_S1_). In S1, we note three types of occurrences: a channel remains closed for the entirety of S2 (C_S2_), it may open and inactivate during S2 (O^I^_S2_) or it may open and remain open for the duration of the depolarization (O^S^_S2_). Similarly, one can assess the likelihood of particular contingencies, e.g., an O*_S1_ event leading either to C_S2_ (O*_S1_→C_S2_) or O_S2_ (O*_S1_→O_S2_). Similarly, C_S1_→O_S2_ and C_S1_→C_S2_ can be assessed. For each of the four channel complexes, Table 1 summarizes event occurrences at the end of segment S1 and during segment S2 (rows 1-7), along with dependencies of S2 behavior on behavior at end of S1 (rows 8-11). From this information, the fractional occurrence of particular S2 behavior (i.e., channel opening (O_S2_) or remaining closed (C_S2_) over the entire data set can be evaluated. Similarly, whether channel status at the end of S1 (O*_S1_ or C_S1_) impacts on likelihood that a channel opens or remains closed during S2 can be evaluated. Although the conformational state (i.e., closed vs. inactivated) of any given closed event can never be known with absolute certainty, we utilize the assumption that a closure persisting for all of S2 most likely represents an inactivated condition. The table includes two columns for WT+D20Am results, one generated with a 150 ms prepulse (column 2) and the other with a 250 ms prepulse (column 3).

**Figure 6.**
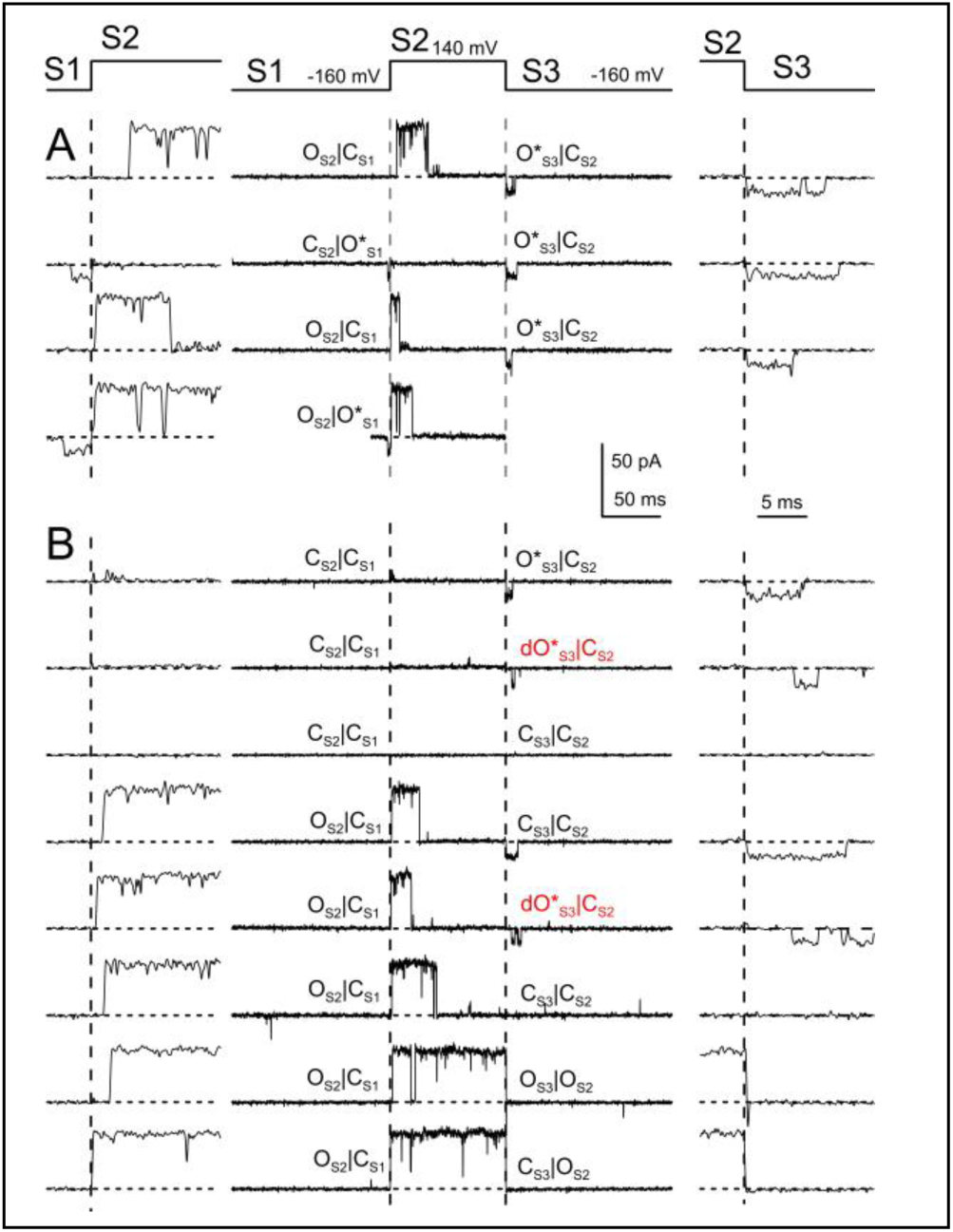
Examples of channel behavior at S1-S2 and S2-S3 voltage transitions. Full voltage-step protocol in highlighted in middle, while trace on left shows behavior at the S1-S2 transition on faster time base, and trace on right highlights S2-S3 transition also on a faster time base**. (A)** Traces illustrate each of 4 possible behaviors at the S1-S2 transition. From top to bottom, C_S1_→O_S2_, O*_S1_→C_S2_, C_S1_→ O_S2_, and O*_S1_→O_S2_. Top three traces are examples from WT+D20Am channels. The bottom one was from RKK/3Q+D20Am and was the only such case observed frmo over 600 O*_S1_ events from all patches. **(B)** Traces (all for WT+D20Am) were selected to highlight the various possible behaviors observed at the S2-S3 transition. From top to bottom, C_S2_→O*_S3_; C_S2_→dO*_S3_; C_S2_→C_S3_; another C_S2_→C_S3_, but with an opening during S2; C_S2_→dO*_S3_, also with an opening during S2; C_S2_→C_S3_ also with an opening during S2; and finally, two rows showing transitions in which a channel is open at the end of S2, one with a partially resolved tail opening, O_S2_→O_S3_, and a case of no detectable tail opening (O_S2_→Cs_3_)

For each construct, the number of occurrences of particular S1 and S2 transition behaviors is given in rows 1-7. Associations between a given S1 event with a subsequent S2 outcome are given in rows 8-11 (Table 1). We note several features of the single channel behaviors for particular attention. First, the likelihood of occurrence of O*-I behavior at the end of S1 substantially increases in the order of WT<3Q<3E<3V (row 3 as fraction of row 1). Second, almost uniformly (except for 1 of 60 O*_S1_ events in RKK3Q), O* behavior at the end of S1 results in the complete absence of any channel opening during S2 (row 13). Thus, occupancy in O* at the end of S1 by definition reflects an inactivated condition.This single channel behavior mirrors the impact of persistent inward current in macroscopic current recordings on subsequent depolarization activated outward current (Fig. 3-4). Third, when a channel is nonconducting at the end of S1 (C_S1_), two distinct outcomes in S2 are observed, given in C_S2_ (row 10) and O_S2_ (row 11). Whereas for WT+D20Am, channels are more likely to undergo a C_S1_→O_S2_ transition, for the three RKK mutants there is increasingly large excess of C_S1_→C_S2_ events. This argues that C_S1_ events are of two types, a channel occupying a closed state that is available for activation and a channel in a closed state that is unavailable to be activated, again by definition an inactivated state, but distinct from O*-I. Thus, this is direct evidence for a long-lived inactivated state that can be observed at negative voltages at the end of the S1 prepulse. The likelihood of occurrence of the long-lived inactivated state at the end of S1 increases in the order of WT <RKK3Q <RKK3E with RKK3V being intermediate between 3Q and 3E.

From the events and associations, we also note the following. Row 12 provides the fraction of all trials in which channel opening occurs during S2: 0.81 (150 ms pp), 0.90 (250 ms pp), 0.35, 0.03, and 0.04, respectively, for WT (two prepulse durations), RKK3Q, RKK3V, and RKK3E. These values match roughly with the fractional peak current amplitude of inactivating macroscopic current normalized to total BK current in patches (Fig. 4B,C). Row 13 summarizes for each construct the fraction of O*_S1_ events that result immediately in inactivation during S2, again highlighting the idea that that occupancy in the O*-I ensemble, despite its measurable current, results in immediate inactivation with depolarization. Row 14 gives an estimate of fraction of sweeps for which channels are in an inactivated state at end of S1, meaning the sum of O*→C_S2_ and C_S1_→C_S2_ over all sweeps. A failure to open during S2 likely reflects channels that are already inactivated at the end of S1. For WT+D20Am channels, this suggests 10-15% of channels are apparently inactivated at the end of S1 in these trials which exceeds the fraction we might have been expected based on our evaluation of fractional reduction of peak macroscopic current based on coupled or uncoupled activation models (as in Fig. 4B,C) where we inferred that very few WT+D20Am channels were likely to be inactivated prior to the depolarizing step. The reasons for this discrepancy are not entirely clear. However, increasing the duration of the prepulse from 150 ms to 250 ms did reduce the fraction of inactivated channels at the end of the prepulse so that may be one contributing factor. Another possibility is that our use of trypsin to define the maximal BK conductance in a patch may result in an underestimate of G_max_ in the macroscopic current, either because of incomplete digestion or because of residual flickery block. Based on the approach used earlier to calculate expectations for model-dependent reductions in peak current (Fig. 4B,C), it can be calculated that an underestimate of BK G_max_ in a patch of about 20% might reconcile the single channel and macroscopic observations. Finally, we note that Row 15 provides the fraction of closures at end of S1 (C_S1_) that can be considered an inactivated-closed, as opposed to a resting closed state. The information in row 15 and 16 makes a strong case that persistent currents alone (O*-I occupancy) are not the only factor contributing to suppression of depolarization-activated outward current in the RKK mutant constructs. Long-lived inactivated states at the end of S1 account for fractions of 0.74 and 0.77 of all C_S2_ inactivation events for RKK3Q and RKK3E after a 150 ms prepulse. In contrast, for RKK3V, 0.90 of C_S2_ inactivation follows O*_S1_ events. Together, these results reveal a longer-lived, non-conducting inactivated state, not considered within the context of the Scheme 1 inactivation model. Additional considerations presented below require that this longer-lived inactivated state does not follow O*-I, but likely represents inactivation occurring from closed states.

### Most inactivated WT and mutant RKK channels recover from inactivation by immediate passage through O*-I behavior

We next evaluated events (Fig. 6B) associated with the S2-S3 transition (Table 2). For all three RRK mutant channels, no trials exhibited an opening at the end of S2 (Table 2, row 1), while for WT there were 36 sweeps in which the channel was open at the end of S2, resulting either in a normal fast deactivating channel tail opening at the beginning of S3 or no resolved opening. For this analysis, we consider trials in which the channel was closed at the end of S2 (C_S2_). Three types of behaviors were defined for S3: 1) cases in which channels immediately entered O*-I burst behavior (O*_S3_), 2) cases in which a channel initially occupied a resolvable non-conducting closure before entering into O*-I behavior with a delay (dO*_S3_), and 3) cases in which no opening was observed during S3 (C_S3_).

**Table 2.** Likelihood of S3 behaviors contingent upon channel status at end of S2.

|  | Event Occurrences | WT (N=5)<br>150 ms pp | WT (N=6)<br>250 ms pp | RKK3Q<br>(N=6) | RKK3E<br>(N=8) | RKK3V<br>(N=8) |
| --- | --- | --- | --- | --- | --- | --- |
|  | Total Trials | 793 | 396 | 359 | 330 | 588 |
|  | O<sub>2</sub>(inactivation) | 608 | 320 | 125 | 10 | 12 |
| 1 | O<sub>S2</sub> (no inactivation) | 37 | 35 | 0 | 0 | 0 |
| 2 | C<sub>S2</sub> (closed at end of S2) | 756 | 361 | 359 | 330 | 588 |
| 3 | C<sub>S3</sub> | 122 | 80 | 0 | 0 | 0 |
| 4 | O*<sub>S3</sub> | 616 | 271 | 347 | 309 | 458 |
| 5 | dO*<sub>S3</sub> | 18 | 10 | 12 | 21 | 17 |
| **Fraction dO\*<sub>S3</sub> / (dO\*+O\*)** |  |  |  |  |  |  |
| 6 | dO*<sub>S3</sub>/(dO*<sub>S3</sub>+O*<sub>S3</sub>) | 0.028 | 0.036 | 0.033 | 0.064 | 0.029 |
| **Fraction of all C<sub>S2</sub> leading to O\*+dO\* events** |  |  |  |  |  |  |
| 7 | (O*<sub>S3</sub>+dO*<sub>S3</sub>) /C<sub>S2</sub> | 0.839 | 0.778 | 1 | 1 | 1 |

O<sub>S2</sub>: sweep with opening at end of S2
C<sub>S2</sub>: channel is closed at end of S2
C<sub>S3</sub>: closures during all of S3.
O\*<sub>S3</sub>: O\* opening immediately after repolarization
dO\*<sub>S3</sub>: O\* opening following a brief closure immediately following repolarization
Row 6. Occurrence of dO\*<sub>S3</sub> behavior is similar among constructs. dO\*<sub>S3</sub> vs. O\*<sub>S3</sub>. Fishers exact with Bonferroni. WT vs 3Q, P=0.1029; WT vs 3E, P>0.9999; 3Q vs. 3E, P=0.6293; WT vs. 3V, P>0.9999; 3Q vs 3V, P=0.9772; 3E vs. 3V, P>0.9999; WT1 vs. WT2, P>0.9999.
Row 7. RKK mutants never close during entirety of S3. O\*<sub>S3</sub> vs. C<sub>S3</sub>. WT vs 3Q, P<0.0007; WT vs 3E, P<0.0007; 3Q vs. 3E, P>0.9999; WT vs. 3V, P<0.0007; 3Q vs 3V, P>0.9999; 3E vs. 3V, P>0.9999; WT1 vs. WT2, P=0.112.

The dO* closures represent an event not imbedded in the simple 2-step inactivation model. The fraction of dO*_S3_ events as a function of all tails with immediate or delayed O*-I behavior is summarized in row 6 (Table 2). This fraction varies from 0.033 for RKK3Q to 0.066 for RKK3V, but, in general, the frequency of occurrence is similar among constructs (statistics at bottom of Table 2). This suggests that during inactivation at positive potentials, about 5% of the time, irrespective of the construct, a channel may occupy a fully non-conducting state distinct from O*-I, but which is likely connected to O*-I. However, the frequency of such events at least as inferred by their occurrence at −160 mV suggests that such events will have small effects on macroscopic tail current amplitudes. To estimate the lifetime of such rare events, for the set of 588 trials with RKK3V+D20Am channels, 17 dO* closures at the beginning of S3 had a mean duration of 19.1±23.6 ms, while at the beginning of S1 (repolarization from 0 mV), 29 dO* closures that preceded O*-I bursts had a similar mean duration of 35.5±36.7 ms (T-test: P=0.114). Given the low frequency of occurrence of this state, such events would reduce the initial peak tail conductance less than 10% from that arising from all channels being in O*-I, so are not considered further here.

For D20Am-associated RKK mutant channels, closures at the end of S2 were invariably associated with either O* or dO* at the onset of S3 (Table 2, row 7). This differs from WT+D20Am channels which fail to open at all during S3 in about 20% of sweeps for which a channel is closed at the end of S2. The high likelihood that inactivated RKK mutant channels immediately enter O*-I bursts upon repolarization suggests that the peak of macroscopic tail current for each of the RKK-mutant channels arises almost exclusively from channels occupying the O*-I state (or dO* state) ensemble at any given repolarization potential. This is important in our considerations below. Furthermore, the macroscopic and single channel observations together argue that the persistent macroscopic inward current for the RKK mutant channels reflects the prolonged fractional occupancy of channels in the O*-I burst behavior typical of the WT+β3a tail current openings. No observation suggests that normal BK openings contribute in any way to such persistent current.

### Inward current of RKK3V+D20Am channels reflects voltage-dependence of O*-I equilibrium

Guided by single channel results indicative that repolarizations from +140 mV to −160 mV are predominantly associated with immediate O*-I openings, we now turn to a closer examination of the macroscopic tail currents following repolarization. The magnitude of the inward tail current at negative potentials arising from each of the D20Am+RKK mutant channels represents a surprisingly large fraction of the total BK conductance that can be activated in a given patch defined following trypsin removal of inactivation. Here we first focus on the persistent RKK3V+D20Am inward current to define the magnitude and voltage-dependence of this conductance. An intriguing aspect of the RKK3V+D20Am current (Fig. 7A, B) is the similarity of the tail current at both 0 and 10 μM Ca^2+^. This suggests that normal Ca^2+^-dependent BK gating transitions (C←→O) are not impacting on the RKK3V+D20Am inward current.

**Figure 7.**
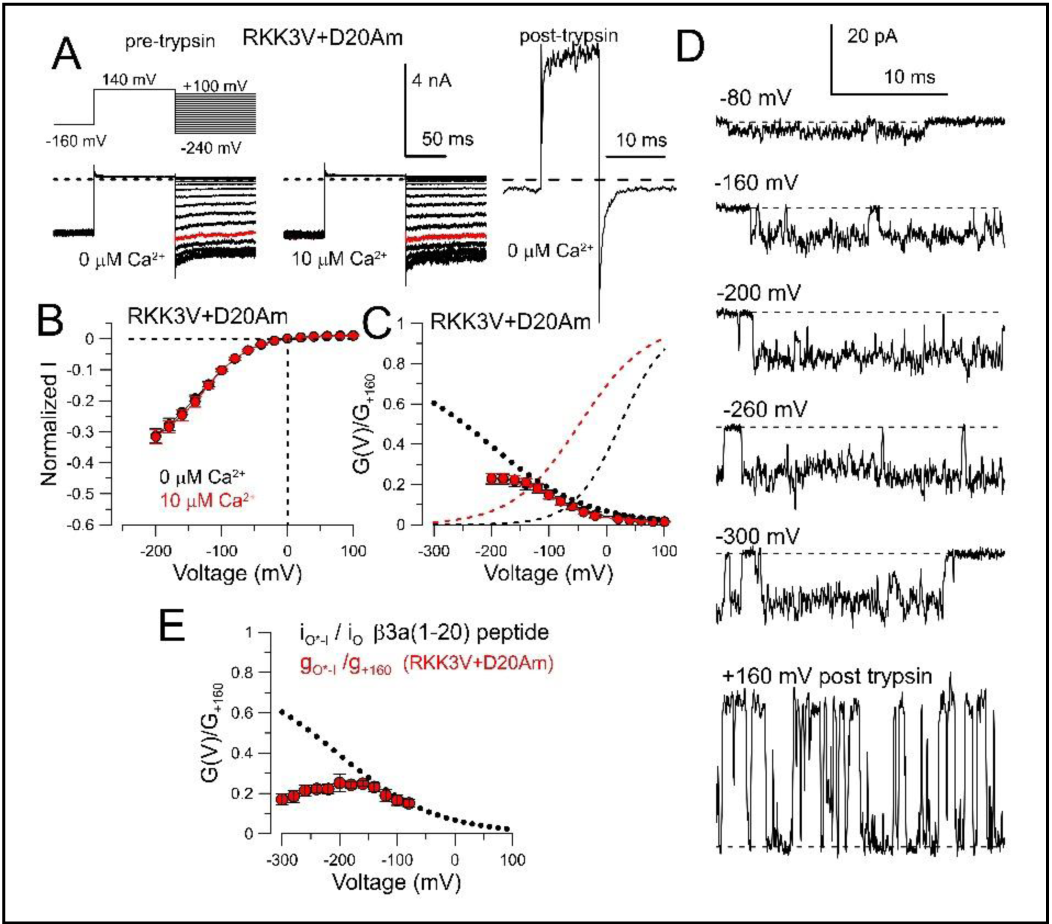
Voltage-dependence of inward current arising from RKK mutant constructs mirrors single channel β3a O*-I equilibrium. A) Examples of persistent inward current from RKK3V/D20AM channels at 0 (left) and 10 (right) μM Ca^2+^. Note complete absence of outward current with depolarization to +140 mV. On the far right, maximum BK current activated at 0 μM Ca^2+^ at +160 mV following trypsin removal of inactivation is shown. All traces from same patch. (B) Voltage-dependence of persistent steady-state current arising from RKK3V/D20Am currents at 0 and 10 μM Ca^2+^ normalized to peak BK current after trypsin. (C) Steady-state current from B was converted to normalized conductance. Dashed lines correspond to activation of RKK3V/D20Am channels after removal of inactivation at either 0 (black) or 10 (red) μM Ca^2+^ with parameters as in Figure 3. Dotted line corresponds to amplitude of WT+β3a(1-20) peptide single channel tail bursts exhibiting O*-I behavior normalized to WT single channel current amplitude (Gonzalez-Perez et al., 2012). (D) Single channel openings of RKK3V+D20Am channels at the indicated voltages, with the trace at +160 mV after removal of inactivation by trypsin. (E) Single channel amplitudes were converted to conductances and normalized to the single channel conductance at +160 mV with the dotted line showing GV curve from the β3a(1-20) peptide inhibition of single WT BK channels (Gonzalez-Perez et al., 2012).

From the average amplitude of steady-state tail current from a set of RKK3V+D20Am patches, we plot normalized IV curves (Fig. 7B) for both the 0 and 10 μM data sets and then GV curves (Fig. 7C). GVs were calculated based on the persistent current amplitudes and the maximum BK current in a given patch from:

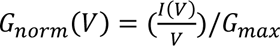

where I(V) is the persistent current amplitude and G_max_ is determined from I_+160_/160 mV following trypsin removal of inactivation. At both 0 and 10 μM Ca^2+^, the net fractional conductance at −200 mV for a patch expressing RKK3V+D20Am is about 0.25 of the maximum BK conductance for each patch (Fig. 7C). The net conductance increases with hyperpolarization, in marked contrast to the voltage-dependence of the normal C←→O activation equilibrium. Overlaid on the GV plot generated from macroscopic persistent current (Fig. 7C), we also plot the activation GV curves measured for RKK3V+D20Am-IR to highlight the idea that there is appreciable and increasing persistent conductance at voltages where activation Po is low.

Based on the idea that the RKK3V+D20Am persistent current arises from occupancy of channels almost entirely in the O*-I ensemble, the voltage-dependence and amplitude of the persistent conductance should reflect the underlying voltage-dependence of the fractional occupancy of a single channel when gating in the O*-I ensemble. Previous work reported the amplitude and voltage-dependence of single channel O*-I bursts resulting from β3a(1-20) peptide inhibition of WT BK channels normalized to the intrinsic single channel current amplitude (Gonzalez-Perez et al., 2012). From the *O*^∗^ ↔ *I* relationship as defined in Scheme 1 (Fig. 2C) and plotted in Figure 2G, if *k_4_* and *k-_4_* reflect the entry and exit rates into I, k_-4_/(k_4_+k_-4_) defines the fractional amplitude of the O*-I burst (normalized to a full open channel current level) as a function of voltage (black dotted line on Fig. 7C). The similarity of the previously measured single channel O*-I behavior to the GV for normalized macroscopic RKK3V+D20Am conductance supports the idea that the persistent inward current and its voltage-dependence also arise from the O*-I behavior. Furthermore, the voltage-dependence of activation of the D20Am+RKK3V persistent current appears intrinsic to the β3a N-terminus and does not seem influenced either by the RKK mutations or any D20Am-induced gating shifts. Qualitatively, the increase in conductance with hyperpolarization is consistent with a reduction in likelihood of blockade by a cytosolic blocking moiety under conditions of pronounced inward current. One difference between the peptide results and the RKK3V+D20Am curves is that the RKK3V+D20Am conductance begins to exhibit a more pronounced roll-over at the most negative potentials. As mentioned earlier, this roll-over likely reflects the inward current rectification that arises from the extracellular loops of the D20Am construct, which are derived from the BK β2 subunit (Zeng et al., 2003), but would not occur with isolated peptide application to WT BK channels. Overall, these results and analysis provide additional support that the persistent current arises from steady-state occupancy in O*-I states.

To test whether the tethered D20Am N-terminus might differ in effects on amplitude of O*-I bursts compared to β3a(1-20) peptide-induced O*-I bursts, we measured amplitudes of RKK3V+D20Am single channels (Fig. 7D) over potentials from −80 to −300 mV and normalized them to BK single channel conductance determined from current amplitude at +160 mV following trypsin and assuming voltage-independence of the conductance. The low Po of RKK3V+D20AM-IR channels precluded acquisition of enough resolved openings following trypsin digestion to assess RKK3V single channel amplitudes at negative voltages. The plot of the fractional O*-I amplitudes of RKK3V+D20Am openings (Fig. 7E) is quite comparable to that of the RKK3V+D20Am persistent conductance, including the expected inward rectification arising from the β2 subunit extracellular loops (Zeng et al., 2003). Thus, the behavior of the O*-I bursts resulting from β3a(1-20) peptide is similar to that of a tethered β3a N-terminus, irrespective of whether inhibition arises from a native WT or RKK3V mutant channel.

### RKK3V+D20Am persistent currents are resistant to paxilline inhibition

The unusual nature of the RKK3V+D20Am persistent current prompted us to test its sensitivity to standard inhibitors of BK current. When RKK3V+D20Am currents were activated with the standard activation protocol at 0 Ca^2+^, 200 nM paxilline failed to produce any inhibition (Fig. 8A1) at voltages from −200 to +140 mV (Fig. 8B). However, after application of trypsin to digest the D20Am N-terminus, paxilline produced essentially complete inhibition (Fig. 8A2, C), consistent with the idea that the persistent currents do arise from BK channels. The failure of paxilline to inhibit the intact RKK3V+D20Am currents is not entirely unexpected, but might arise from either of two reasons. First, occupancy of the RKK3V central cavity by D20Am may maintain the BK conformation in an open state. In earlier work (Zhou and Lingle, 2014) paxilline has been shown to inhibit the closed channel and be unable to reach its likely blocking position in the open state (Zhou et al., 2020). Second, binding of the D20Am N-terminus may also sterically prevent paxilline from reaching its site of inhibition. We found that, in contrast to paxilline, 20 mM TBA did produce a modest inhibition of RKK3V+D20Am persistent inward current (Fig. 8D1, E), while after removal of inactivation 20 mM TBA produce profound inhibition of the outward BK current (Fig. 8D,F). The modest TBA inhibition of intact RKK3V+D20Am channels is unlikely to be accounted for solely by the known voltage-dependence of block (*z*=∼-0.15e (Li and Aldrich, 2004)), but probably arises from a somewhat weaker TBA apparent binding affinity for the O* state compared to the O state. This is consistent with previous observations showing that TBA speeds up the slow tail current following repolarization, consistent with some inhibition of the O* state (Gonzalez-Perez et al., 2012). Together these results provide further support for the idea that the D20Am N-terminus occupies a position in the BK central cavity that may influence access of standard BK inhibitors to their binding sites.

**Figure 8.**
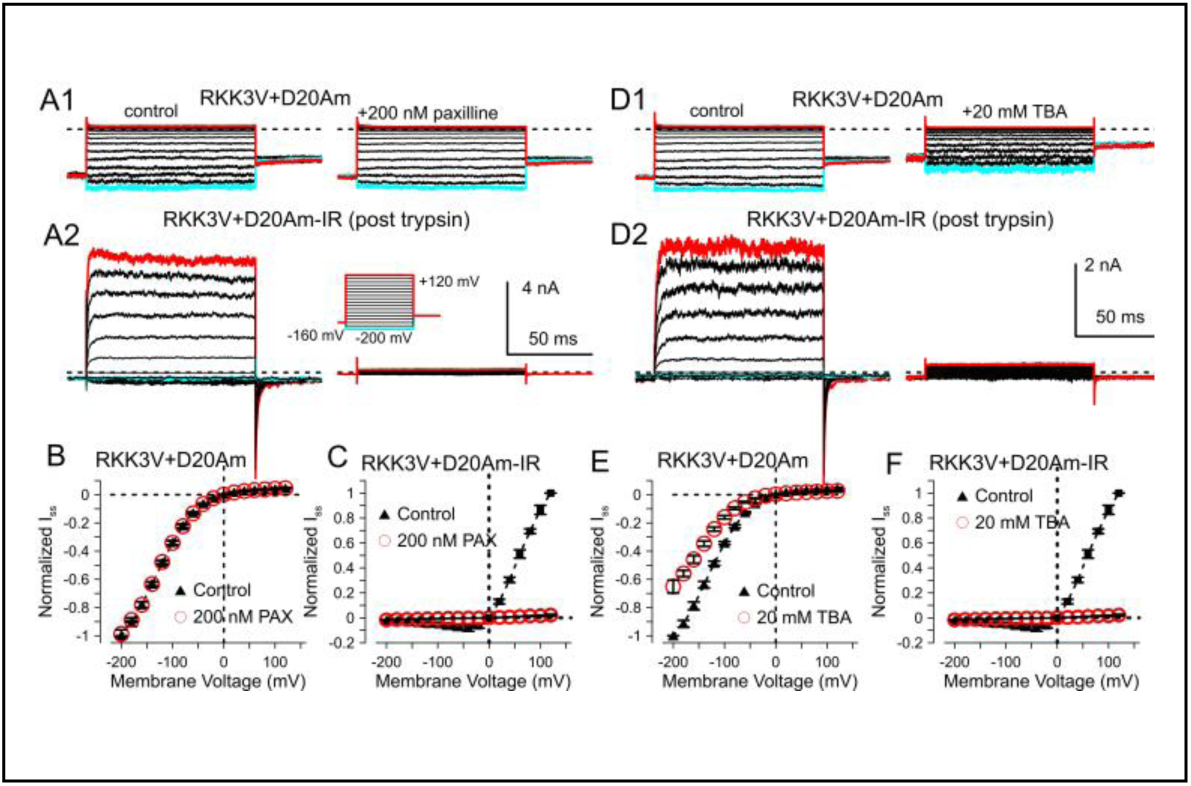
RKK3V+D20Am currents are insensitive to paxilline. A1) Left, RKK3V+D20Am currents were activated with a standard activation protocol with voltage-steps from −200 to +140 mV with normal symmetrical K_o_/K_i_ gradients. Right, 200 nM paxilline was applied. Red trace: +140 mV; blue trace: −200 mV. A2) Left, currents from the patch shown in A1 following trypsin application. Right, 200 nM paxilline was applied. B) Steady-state current normalized to the value at −200 mV is plotted for different activation voltages without and with 200 nM paxilline, from traces as in A1. N=5 patches. C) IV plots for steady state currents after treatment with trypsin without and with 200 nM paxilline. D1) Example RKK3V+D20Am currents from a different patch before (left) and after (right) application of 20 mM TBA. D2) Currents after application of trypsin before (left) and (after) application of 20 mM TBA. E) IV curves for steady-state RKK3V+D20Am currents without and with 20 mM TBA. F) IV curves after trypsin treatment without and with TBA.

### State occupancies at beginning and end of recovery from inactivation

In order to further tease apart the impact of the RKK mutants on D20Am inactivation, macroscopic tail currents at both 0 and 10 μM Ca^2+^ were elicited by repolarization from +160 mV to voltages from −240 to −80 mV for all four constructs (WT+D20Am (Fig. 9A), 3Q+D20Am (Fig. 9B), 3E+D20Am (Fig. 9C), and 3V+D20Am (Fig. 9D). For WT, RKK3Q, and RKK3E, peak tail amplitude was somewhat larger at 10 μM Ca^2+^ than with 0 Ca^2+^, while tail current decay was slowed, consistent with the idea that at higher Ca^2+^ more channels are inactivated at the time of repolarization and that, as some channels return to resting states, they may be reactivated and inactivated, thereby slowing the tail current. In contrast, RKK3V tail currents were markedly similar at both 0 and 10 μM Ca^2+^, suggesting that Ca^2+^-dependent gating transitions have little impact on state occupancies during the RKK3V persistent tail currents. For WT, RKK3Q, and RKK3E, tail currents were fit with both single and double exponential decay functions, with the two exponential fit providing the best description of the decay process with time constants and fractional amplitudes summarized in Figure S2. We note the following features of the tail currents. First, tail current decay becomes faster at more negative voltages for WT, RKK3Q, and RKK3E. Second, based on the extrapolated baseline of the two exponential fits, there appears to be persistent inward current for both RKK3Q and RKK3E even at voltages from −160 to −240 mV, although in each case more than half the channels do appear to return to non-conducting states. Third, for RKK3V, the lack of Ca^2+^-dependence of the tail currents and the persistent nature of the currents suggests that RKK3V+D20Am channels may be essentially locked in O*-I gating behavior at least for the repolarization durations we have tested.

**Figure 9.**
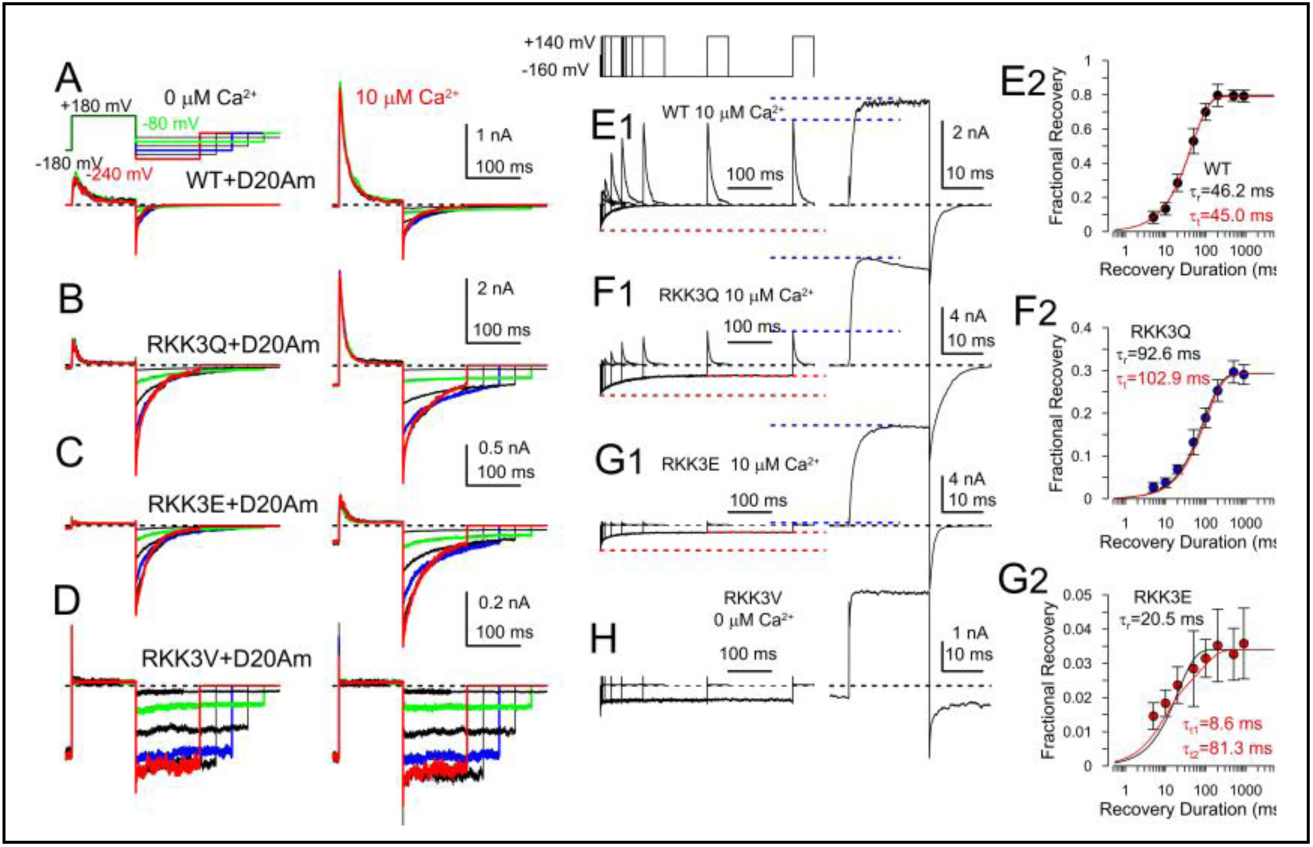
Voltage-dependence of tail current relaxations, persistent current, and time course of recovery from inactivation. (A) The indicated voltage protocol was used to examine variation in tail current properties following complete inactivation at +180 mV of WT+D20Am channels at either 0 Ca^2+^ (left) or 10 μM Ca^2+^ (right), both from same patch. Tail current voltage duration was shorter for more negative voltages to help maintain patch viability. Tail currents at −80 mV, −160 mV, or −240 mV are highlighted in green, blue, and red, respectively. (B) Currents for RKK3Q+D20Am channels. (C) RKK3E+D20Am currents. (D) RKK3V+D20AM currents. (E1) The paired pulse protocol on the top was used to monitor recovery from inactivation of outward current at −160 mV with the indicated cytosolic [Ca^2+^] for WT+D20Am channels. The trace on the right defines total BK current in the patch following removal of inactivation by trypsin, with the same vertical calibration both before and after trypsin. Dotted red lines highlight the peak tail current level and the steady-state level of the tail current which was essentially 0 for WT. Blue lines highlight peak inactivating current amplitude following full recovery and the amplitude with inactivation removed. (E2) Points plot time course of recovery of peak outward current normalized to total BK current (N=4 patches). Red line represents the normalized slow component of the two exponential fit to the tail current resulting from the 900 ms recovery step. (F1) Time course of recovery for RKK3Q+D20Am currents, along with trace (right) after trypsin digestion. Dotted lines are peak and steady-state levels determined from a two component exponential fit to the decay. (F2) Plot of time course of recovery (N= 5) of peak amplitude as in panel F1, along with fit of the slow exponential component of the tail during 900 ms recovery step at −160 mV. (G1) RKK3E+D20Am channels, along with trypsin-digested currents at +160 mV (right trace). Dotted lines are as in panel F1. Note that, although about 70% of channels return to closed state during the tail current relaxation, only about 3% of channels activate during the subsequent depolarization. (G2) Plot of recovery of peak outward current for RKK3E+D20Am currents (N=5) normalized to amplitude after trypsin digestion along with fit of one and two component exponentials to tail during 900 ms repolarization step. (H) Example of currents for RKK3V+D20Am using the same paired pulse recovery protocol. No recovery in outward current was measurable.

A trivial expectation for the D20Am-mediated tail currents is that the time course of the tail currents should match the time course in recovery of channels that become available to be activated with depolarization. We therefore used a paired pulse protocol to check this expectation (Fig. 9E1, F1,G1,H). After 200 ms at 0 mV to produce inactivation, the duration of the repolarizing step to −160 mV was varied between 5 ms up to 900 ms followed by a depolarizing test step to +140 mV to monitor the fraction of channels that recover from inactivation in each case. At the end, trypsin application defined the maximum BK current in the patch allowing the fractional recovery for each construct to be plotted in terms of maximal possible BK current (Figs. 9E2,F2,G2). Then, for the trace with the 900 ms repolarization, the slow component of the tail relaxation was measured. The peak and steady-state levels of the fit to the tail currents were then normalized to the time course of outward current recovery for WT (Fig. 9E2), RKK3Q (Fig. 9F2), and RKK3E (Fig. 9G2). Given the limited recovery durations that were measured, any fast component of recovery in outward current would not be resolved. However, for these three constructs, the slow component of tail relaxation matched well with the recovery time course in outward current amplitude. In contrast, RKK3V+D20Am did not show any obvious outward current at +140 mV and only minimal if any relaxation in the tail current at −160 mV (Fig. 9H). Overall, for WT, RKK3Q, and RKK3E, the time courses of recovery in paired pulse amplitude match well with the time courses of the respective slow component of tail current decay (Fig. 9E1,F1,G1). For each of the RKK mutant constructs, recovery from inactivation appears to have reached a near steady-state level at −160 mV within about 200 ms, suggesting that, at potentials of −160 mV and more negative, prepulses of 200 ms should be near an equilibrium distribution among states.

Despite the return of WT, 3Q, and 3E channels to closed states that permit activation, inspection of the amplitude of voltage-activated outward current during the paired pulse protocol relative to fractional decay of the tail current reveals an intriguing discrepancy. The decay of peak tail current to steady-state tail current is highlighted with dotted red lines (Fig. 9E1-7G1). The difference between the peak and steady-state during the tail current relaxations can be considered to reflect the fraction of channels that have returned to closed states, and therefore might be expected to be available for activation. The dotted blue lines highlight the difference between the channels that were available for activation, as indicated by the peak outward current prior to trypsin compared to the total number of channels following removal of inactivation. For WT channels, the example tail currents suggest almost all channels return to closed states, which is consistent with the voltage-activated outward current being about 80% of the maximal BK conductance, after correction for the 15-20% reduction expected for inactivation occurring during the rising phase and impact of asynchrony in opening (Fig. 4B). In contrast, for 3Q and 3E, although the tail currents suggest that 64.5% and 71.0% of channels may have returned to rest, the peak outward current after recovery indicates that only about 31.6% and 3% of channels have really returned to closed states. This suggests that some fraction of the channels that return to closed states during the tail currents and are no longer contributing to O*-I persistent current are not available for activation. The implication is that this fraction of channels are closed and inactivated. This implication is congruent with the long duration inactivated states inferred from analysis of closed states preceding the S1-S2 transition in single channel records (Fig. 5) and also is consistent with the steady-state inactivation curves (Fig.4D).

To address this observation that fraction of channels that have returned to closed states greatly exceeds the fraction that become available for activation, we examined the voltage-dependence of the tail current relaxations (as in Fig. 9) following careful subtraction of capacity transients. As shown for example tail currents at −240, −200, and −160 mV for WT (Fig. 10A), 3Q (Fig 10B), 3E (Fig. 10C), and 3V (Fig. 10D), we measured peak tail amplitude (I_Inst_) and fit 1 and 2 exponential functions to the decay time course of the tail current. We used the steady-state value from the two exponential fit to the current decay as a measure of the steady-state persistent current (I_ss_) (Fig. 10A-C). For each construct, the peak tail current at each voltage was converted to a fractional conductance based on the maximum current at +160 mV after trypsin (Fig. 10E). Despite some differences among constructs in average G(V)/G_max_ (Fig 10E), the peak tail current amplitudes after conversion to conductances for all four constructs generally track (O*+I)/O relationship defined previously from β3a(1-20) peptide effects on average single channel conductance (Gonzalez-Perez et al., 2012). This again highlights the idea that the tail current peak following repolarization after complete inactivation reflects occupancy of most channels in the patch in the O*-I ensemble. One might expect that the initial peak of the WT and RKK3E tail current would be somewhat less than the others because their faster initial tail current decay (Fig. S2) may result in an underestimate of the apparent peak amplitude. Furthermore, the WT+D20Am tail currents would also be expected to be reduced in amplitude, because of the 20% of WT+D20Am single channels that never open following repolarization.

**Figure 10.**
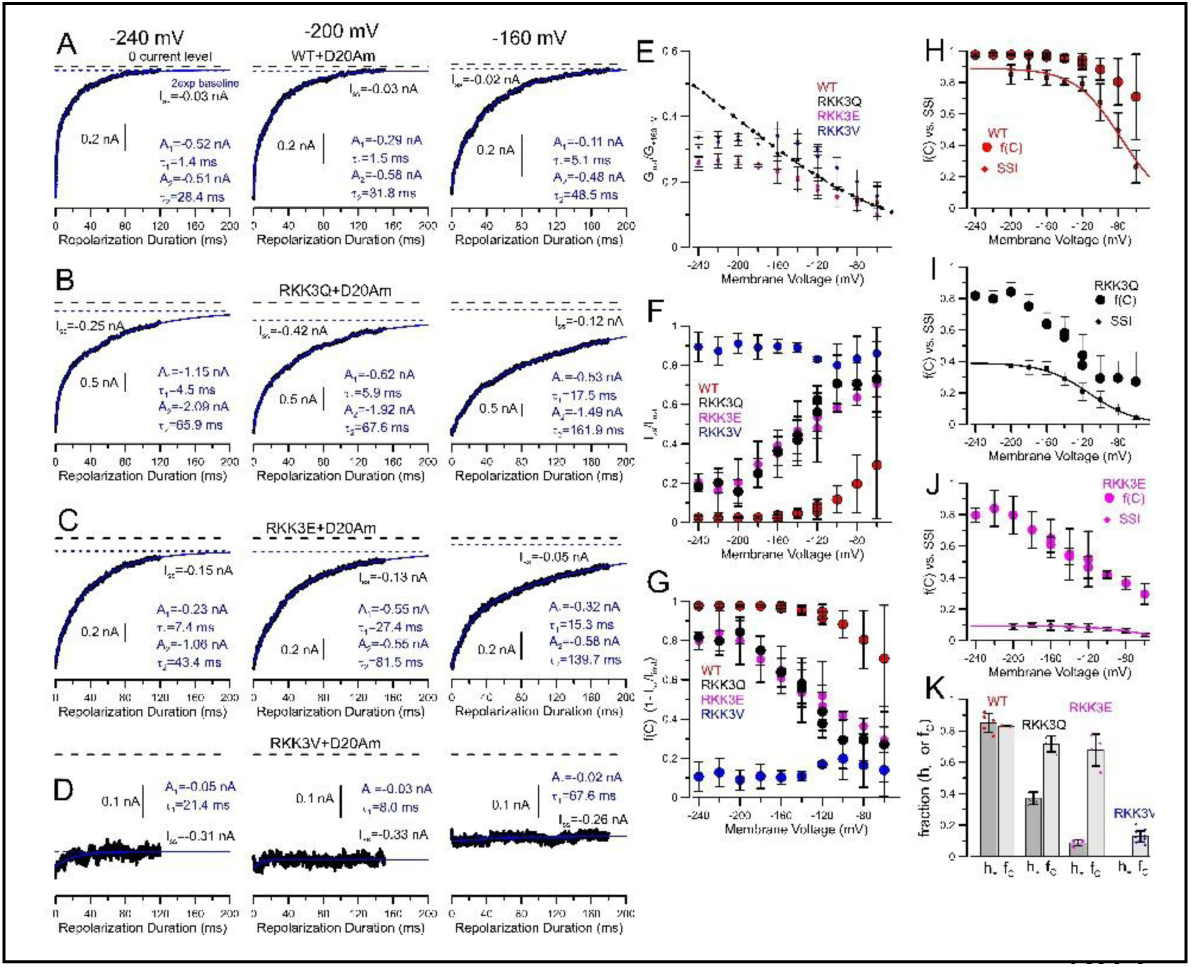
The fraction of inactivated channels that return to closed states is much larger than the fraction that become available for activation. (A) Cm-corrected tail currents for WT+D20Am current (with 10 μM Ca^2+^) at −240 mV (left), −200 mV (center), and −160 mV (right) were fit with a two (blue) exponential function with the dotted lines highlighting the zero current level (black), and baseline of the two exponential (blue) fit. The peak tail amplitude was used to determine tail conductance normalized to that at +160 mV. (B) Similar records for RKK3Q+D20Am reveal a larger component of steady-state current at all voltages. (C) Similar records for RKK3E+D20Am also show a more substantial persistent inward component. (D) Records for an RKK3V+D20Am patch along with single exponential fits with blue line highlighting I_ss_. (E) For a set of patches for each construct, the peak tail current was converted to a relative conductance based on G_max_ determined at +160 mV and plotted as a function of voltage. Each construct results in similar fractional activation at each voltage, consistent with the instantaneous tail current reflecting exclusively the properties of the O*-I equilibrium at each voltage. Dotted line corresponds to the fractional O*-I/O voltage dependence as in Fig. 6C. (F) The fraction of steady-state tail current normalized to peak tail current was determined for each construct. (G) A replot of panel E as 1-(I_ss_/I_inst_) defining fraction of channels that have returned to closed states. (H-J) Comparisons of steady-state inactivation curves for WT+D20Am (H), RKK3Q+D20Am (I), RKK3E+D20Am (J) currents (Fig. 4) to fraction of channels for each construct in closed resting plus closed-inactivated states (f_c_) (panels F,G). (K) Mean and individual h∞ (for WT, 3Q, 3E, N=5,5, and 4, respectively) and f_C_ (for WT, 3Q, 3E, and 3V, N=4, 4, 4, and 8, respectively) values. Using ANOVA with Tukey’s multiple comparison to compare h_∞_ vs. f(C), for WT, P=0.504, for RKK3Q, P=7.13E-06, and for RKK3E, P=2.82E-05.

Qualitatively, the steady-state level of tail current at −240, −200, and −160 mV (Fig. 10A-C) indicates that a large fraction of channels that contribute to peak tail current do return to non-conducting states. For each construct, the ratio of I_ss_/I_inst_ was determined for each patch and plotted over voltages from −60 to −240 mV (Fig. 10F). At the most negative voltages, this ratio essentially defines the fraction of channels still contributing to persistent current. Plotting 1-I_ss_/I_inst_ (Fig 10G) provides an estimate of the fraction of channels that have returned to non-conducting states (fraction of closed channels: f(C)) at the end of the steady-state level of tail current. Expressed this way, essentially all WT+D20Am channels return to closed states over the range of voltages from −120 to −240 mV. In contrast, a fraction of about 0.8 of RKK3Q and RRK3E return to closed states. When applied to RKK3V channels, the average current difference between the beginning of the tail repolarization to that at the end suggests that perhaps 10% of RKK3V channels return to closed states, but this estimate is based on rather uncertain relaxations for RKK3V (Fig. 10D). The most likely error in this approach of measuring the fraction of channels that have returned to closed states would be uncertainty in the extrapolated steady-state current level from the two exponential fit. However, visual inspection of current decays and the extrapolated steady-state currents, particularly at potentials of −200 mV and more negative, suggests this potential error is unlikely to account for the differences calculated between available channels and closed channels.

If all channels that return to closed states are then available for activation, one would expect that the voltage-dependence of f(C) should be similar to the fractional availability curves determined earlier (Fig. 4D). For WT+D20Am channels, the steady-state inactivation curve is quite similar to f(C) (Fig. 10H) if one corrects the f(C) curve to include the 10-20% reduction of peak outward current. At potentials of −100 mV and more positive, the f(C) curves may also be influenced by some fraction of channels that occupy open states (O), so some deviation between the two curves might be expected. For RKK3Q+D20Am channels, the limiting asymptotes at −240 mV for the steady-state fractional availability curves and f(C) differ markedly (Fig. 10I), supporting the idea that, at negative potentials, a large fraction of D20Am-associated channels occupy closed states unavailable for activation, while the difference is even more pronounced for D20Am-associated RKK3E channels (Fig. 10J). By definition, closed channels which are unavailable for activation must be inactivated states, and under the conditions of these experiments these represent inactivated states in equilibrium with resting closed states. Comparison of mean values and individual estimates for h∞ for the set of constructs (Fig. 10K) supports the idea that, although for WT channels inactivation from closed states is either rare or absent, for RKK3Q and RKK3E closed-state inactivation is pronounced.

The situation for RKK3V+D20Am seems more complicated to rationalize. The macroscopic protocols suggest that exit from the O*-I ensemble of states for RKK3V channels to non-conducting states is rare, although the set of single channel patches did identify long-lived closures at the end of the S1 prepulse that was followed by constant closure during S2 in 20% of trials. Based on that, one might have expected that the RKK3V tail current at −160 mV might have exhibited a relaxation of perhaps about 20% from the initial current following repolarization. Although it seems that RKK3V+D20Am channels might be considered locked in O*-I states, perhaps with longer periods at negative potentials such channels might return to closed states.

Overall, this evaluation of f(C) and the comparison to the steady-state inactivation curves (Fig. 4D) provides additional support for the idea that, in addition to some fraction of channels that remain in the O*-I ensemble behavior during the tail currents, there is an appreciable fraction of closed channels that are unavailable for activation, presumably with a β3a N-terminus bound within the central cavity in a position of inactivation.

### A simple, linear 2-step inactivation model explains many D20Am effects on WT and RKK-mutant channels, but not all

An assessment of the D20Am effects within the context of a full BK channel gating scheme (Horrigan and Aldrich, 2002) is beyond the scope of this work. However, here we ask whether major features of the results can be approximated by simple models, that may provide insight into key mechanistic details of D20Am action.

Beginning with the 5-state 2-step inactivation model (Fig. 2C), we constrained most of the defined rates based on experimental considerations. Scheme 1 utilizes a 3-state activation scheme (C1←→C2←→O) and the corresponding rates were adjusted such that the model-predicted GV curves and the time constants for activation and deactivation closely match the experimentally measured properties for WT+D20Am-IR currents (Fig. S1). Since the onset of inactivation of WT+D20Am current is about 16 ms, we set the O→O* transition to 60/s. For the O*-I equilibrium, we set initial rates for the transitions in the O*←→I step (*k*_4_ and *k*_-4_) to be those defined by the earlier evaluation of the single channel burst amplitude with β3a(1-20) peptide (Gonzalez-Perez et al., 2012). The voltage-dependence of this O*/I equilibrium was shown above to be consistent with the voltage-dependence of the amplitude of RKK3V+D20Am single channel amplitude (Fig. 6D-E). The estimates of the absolute 0-voltage rates are undoubtedly not well-defined, but we chose previously described values (Gonzalez-Perez et al., 2012). We did not attempt to incorporate the inward current rectification behavior produced by the β2 subunit extracellular domain into the simulations. Thus, all rates denoted in Scheme 1 (Fig. 2C; Table 3) are guided by experimental observations, except for the O*→O transition (*k*_-3_), i.e., the return from the preinactivated state to the open state, for which we have no direct estimate. As illustrated below, adjustment of this one transition strongly influences the properties of simulated currents and appears likely to be the major contributor to key aspects of the effects of D20Am on WT and RKK-mutant channels. For reasons developed below related to closed channel block, we assigned no voltage-dependence to the O←→O* transitions.

**Table 3.**
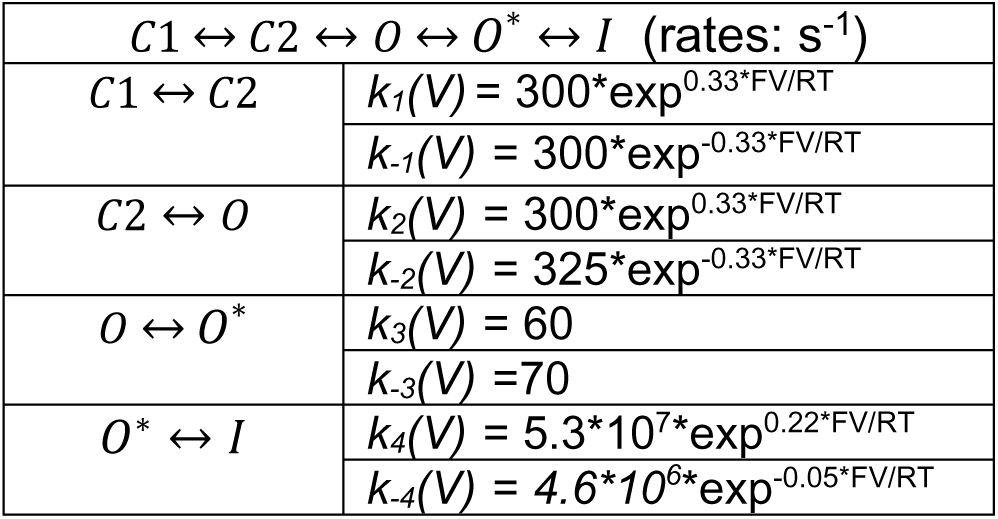
Basic rates used for simulations.

With the above rates, we used three protocols identical to those used in our recordings of D20Am-containing BK currents, first, an activation protocol, second, a tail current protocol, and, third, a steady-state inactivation protocol (Fig. 11A,B). In each case, the initial occupancies among the 5 states at the time of initiation of a given protocol was defined by the steady-state distribution at 0 mV. This results in occupancies preceding any initial negative prepulse defined primarily by equilibration between O* and I. Figure 11A (left to right) presents representative examples of experimental WT+D20Am currents for each of the stimulation protocols while Figure 11B (left to right) displays simulated currents. The simulated currents reasonably capture the fractional reduction in peak current, and the slow tail currents that follow inactivation and are removed by trypsin. Although the tail currents at −120 mV in the activation protocols (Fig. 11A,B left) are of comparable magnitude between experimental and simulated currents (Fig 11A,B left traces), tail currents examined over a range of deactivation potentials protocol(Fig. 11A,B middle traces) exhibit larger tail current amplitude for the simulated currents at more negative voltages. This difference in part likely reflects the rectification behavior of the native WT+D20Am channels. The simulated tail currents also fail to capture a faster component of the tail current that is observed in experimental records.

**Figure 11.**
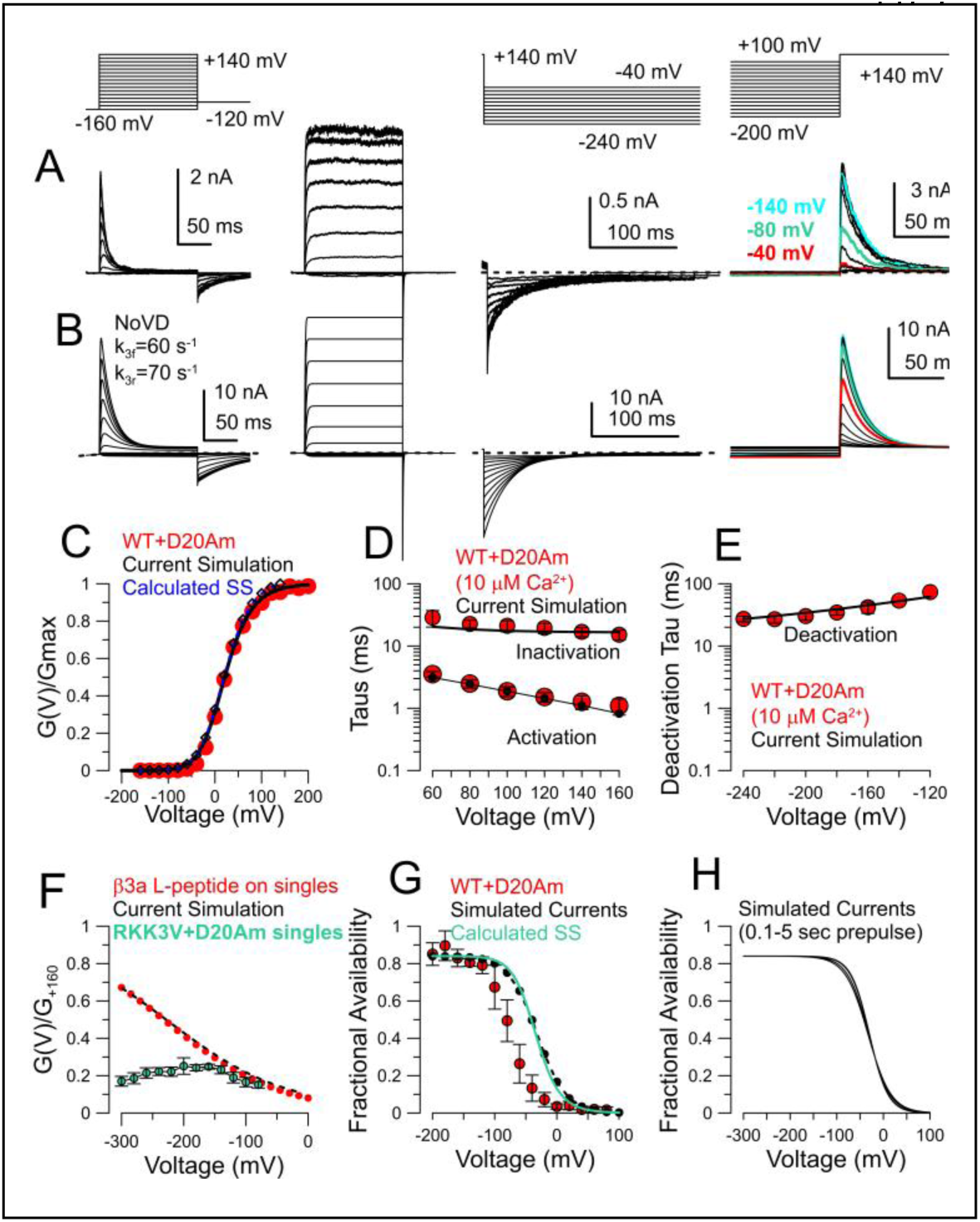
A simple 5-state model incorporating two activation steps and two step-inactivation recapitulates many features of WT+D20Am behavior. **(A)** WT+D20Am currents were evoked by activation (left), deactivation (middle), and steady-state inactivation (SSI) (right) protocols, all with 10 μM Ca^2+^. Currents on left are shown both before and after removal of inactivation by trypsin. Currents evoked by activation and deactivation protocols were from the same patch, and the SSI currents from a different patch. (B) Currents were simulated with same protocols using rate constants in Table 4. Relative amplitude of simulated tail currents greatly exceeds recorded tail currents in part because of marked current rectification, arising from β2 subunit extracellular loops. (C) Measured WT+D20Am-IR GVs (red symbols) are compared to GVs from simulated currents with inactivation removed (D) Measured WT+D20Am inactivation and activation time constants are compared to measured time constants from simulated currents). (E) Experimentally measured and simulated slow time constants of deactivation. (F) Voltage-dependence of simulated instantaneous tail current (black triangles) compared to previous measures of G(V)/G_max_ for fractional reduction of BK single channel current amplitude by β3a(1-20) peptide (red) and RKK3V+D20Am single channel conductance (green) (G) Experimentally measured SSI curves for WT+D20Am (red, example traces on far right of panel A) compared to simulated (black) and calculated (green) SSI curves based on Table 4 rates. (H) Currents from a steady-state inactivation protocol were simulated with prepulse durations spanning 100 to 5000 ms.

GV curves were generated from the simulated currents of scheme 1 and matched well with the average GVs from WT+D20Am-IR currents (Fig. 11C). Similarly, activation and inactivation time constants from simulated currents were similar to experimental values (Fig. 11D), although there appears to be modest voltage-dependence in the experimental inactivation time constants, whereas we assigned no voltage-dependence to the k_3_/k_-3_ rates. The simulated and experimentally measured slow components of deactivation were also comparable (Fig. 11E). The fractional activation of the simulated peak tail conductance tracked that previously determined for β3a(1-20) peptide effects on BK single channel amplitude as expected based on our assumptions (Table 4), but, as expected, deviated from the fractional conductance of RKK3V+D20Am single channel openings which exhibit marked inward current rectification (Fig. 11F). Finally, the experimentally measured steady-state inactivation curve for WT+D20Am was compared to the simulated SSI curve (Fig. 11G). In this case, the model does not adequately account for the position of the SSI curves on the voltage-axis. This discrepancy is not a function of the duration of the prepulse used for the simulated steady-state inactivation protocol (Fig 10H). However, overall, despite some discrepancies between data and model in regards to a faster component of tail current and position of SSI curves, the simple 5 state model does a reasonable job of approximating many features of the experimentally currents. Importantly, unique to the 2-step inactivation portion of the model, the tail currents exhibit instantaneous reopening on repolarization and a net current flux that greatly exceeds that expected for simple, open channel block (Armstrong, 1971; Neher and Steinbach, 1978; Demo and Yellen, 1991).

Having defined a set of rates that approximate many of the key features of WT+D20Am currents elicited with the protocols we used, we next considered whether the major impact of the RKK mutations might be explained largely by an effect on one specific rate in Scheme 1. The relative amplitude of the initial tail current amplitude and the persistent tail currents suggest that the O*/I equilibrium and rates k_4_ and k_-4_ are largely similar for different RKK mutants. Onset of inactivation was not markedly different in cases where it could be measured (3Q,3E). We therefore asked whether changes in *k*_-3_ (corresponding to the N-terminus unbinding, O*→O) might mimic the behavior of the RKK mutant channels. Currents were simulated with k_-3_ values of 100, 70, 30, 10, 2, and 0.5/s (Fig. 12A, top to bottom). Slowing k_-3_ progressively reduced the voltage-activated peak inactivating current, slowed the tail current, and, for the durations of the protocols used, generated persistent inward current (Fig. 12A), capturing each of the notable features of the D20Am-mediated effects on the D20Am+RKK mutant channels. Manipulations of rates, k_4_ and k_-4_, although having effects on the voltage-dependence of initial instantaneous tail current, did not on its own impact on the occurrence of reduction of peak outward current, tail current decay or persistent inward current. Thus, in large measure, these simple simulations suggest that a major effect of the RKK mutations may be to differentially influence the rate of dissociation of the N-terminus from its binding site in the BK central cavity.

**Figure 12.**
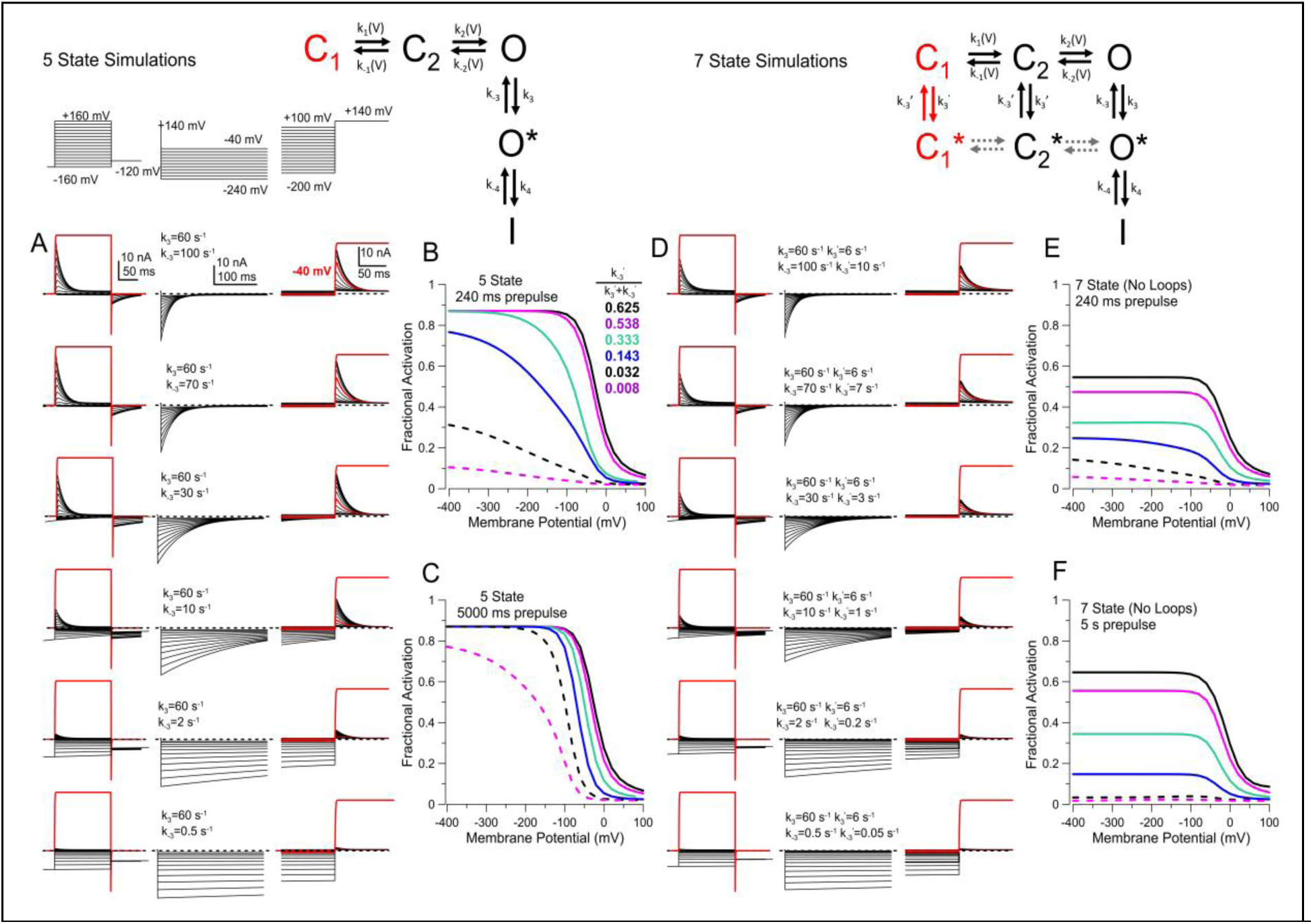
Changes in rate of O*→O mimic aspects of D20Am+RKK mutant currents, but inclusion of closed state inactivation is necessary to explain limiting hinf values in fractional availability curves. (A) Top to bottom, k_-3_ was varied from 100, 70, 30, 10, 2, and 0.5/s and currents simulated with the 5 state model of Scheme 1 with protocols shown at the top. (B) Fractional availability curves were generated from traces on the right side of A, with a 240 ms prepulse. (C) Fractional availability curves from currents generated with a 5000 ms prepulse showing that Scheme 1 predicts identical ℎ_∞_ values in all cases. (D) Similar simulations with the 7 state model with rates (k_3_’=k_3_”, k_-3_’=k_-3_”) for C←→C* transitions given on the panels. (E) Fractional availability with 240 ms prepulses is shown. (F) Fractional availability with 5000 ms prepulses.

From the simulated steady-state inactivation currents, fractional availability curves were then generated (Fig. 12B) based on a 240 ms prepulse duration. This predicts a leftward shift in fractional availability consistent with slowed recovery from inactivation at negative potentials as a function of slower *k_-3_*. However, when steady-state inactivation curves were determined from simulated currents with a 5000 ms prepulse (Fig. 12C), essentially all channels are then able to return to closed states, eventually reaching the same limiting fractional availability. Thus, as argued above, the 5-state model fails to describe steady-state inactivation behavior. With the 5 state scheme, fractional availability curves will always result in the same limiting asymptote, as all channels occupy the most resting closed state, C_1_, at sufficiently negative potentials.

### Closed state inactivation can account for different limiting asymptotes in fractional availability curves

Guided by the experimental results, the 5-state scheme was expanded to include closed-state inactivation (Scheme 2; Fig. 12D, top). Here we only consider expectations for a model in which transitions among C_1_*←→C_2_*←→ O* are rare, as symbolized by the arrows with dotted lines in Scheme 2 (Fig. 12D). The impact of inclusion of transitions among C_1_*, C_2_*, and O* states is addressed in Discussion. Currents were simulated with Scheme 2 utilizing exactly the same rates as given in Table 3, while then adjusting rates of entry and exit from C* states. *A priori* we have no direct information to guide choices for rates of entry into C_1_* or C_2_*, so the outcome of these tests can be considered a bit *ad hoc*. However, from the perspective that the experimental results require that transitions to closed states are occurring, these tests place some constraints on properties of the C_1_/C_1_* and C_2_/C_2_* equilibria. A first consideration is that, irrespective of exact rates, the ratio, k_-3_^”^/(k_-3_^”^+k_3_”), will define the fraction of channels that will occupy C_1_ at the most negative voltages, and are therefore available for activation. Thus, in essence this ratio reflects the limiting ℎ_∞_ asymptote of fractional availability curves.

Interestingly, if one calculates that ratio based on the O←→O* transitions, for k_-3_/(k_-3_+k_3_) one obtains a value of 0.538. This would be the expectation for ℎ_∞_ of the fractional availability curve, uncorrected for inactivation during the rising phase of current activation, if the D20Am N-terminus bound to closed WT channels with an affinity identical to open WT channels. That number is not too dissimilar to the ℎ_∞_value of 0.4 measured in Fig. 4D for D20Am+RKK3Q currents. This suggests that, for D20Am+RKK3Q, the C_1_/C_1_* equilibrium is only somewhat different from that of the WT O/O* equilibrium. A second consideration for the 7-state simulations is that the fact a voltage-independent ℎ_∞_ asymptote is reached requires that the C_1_/C_1_* equilibrium must be voltage-independent. Although this does not necessarily demand that N-terminal binding to the open state is also voltage-independent, that is the reason we chose a voltage-independent O/O* rather than assigning modest voltage-dependence to those underlying rates.

Based on these considerations, we explored various values for k_3_”, k_-3_”, k_3_’ and k_-3_’. When rates from C_1_*→C_1_ and C_2_*→C_2_ are comparable to that for O*→O, say, 60/s, upon depolarization, this results in an additional slow transitions from C_1_* and C_2_* into the activation pathway. This produces a slowing of entry of an appreciable fraction of channels into O, impacting not only on rise time of outward current, but also the inactivation time course. This is contrary to observations and suggests that the underlying C←→C* transitions are likely to be much slower than those (O←→O*) when the channels are open. On the other hand, if rates of entry and exit of N-termini from closed channels are too slow, this essentially removes an appreciabe fraction of channels from ever opening during the protocols we have used. Furthermore, one would then expect appreciable long-lived inactivated states immediately following repolarization, something which is not observed for any of the RKK mutants. Therefore, after exploring various values, to illustrate the impact of inclusion of closed-state inactivation on simulated currents, we chose rates of entry and exit from closed states that were simply 10-fold less than the rates in and out of open states. The resulting simulations for current activation, deactivation, and steady-state inactivation (Fig. 12D) retain the key features of reduction in outward current, slowing of tail currents, and persistent inward current. However, not surprisingly, they support the idea that differences in rate of unbinding of the N-terminus from closed channels underlies the differential ℎ_∞_ values for the different RKK mutants, whether observed with 240 ms (Fig. 12E) or 5000 ms (Fig. 12F) prepulses.

These considerations provide additional mechanistic justification for the idea that the RKK mutant BK channels permit binding of the D20Am N-terminal in closed channels and help explain how modulation of ℎ_∞_ values might occur. However, these simulations are not meant to imply that all the nuances of this inactivation mechanism are now clear. For example, there remains fast components of the tail currents that are not accounted for by the simulations. An attempt to place the inactivation behavior described here within a more realistic BK current activation model might help account for additional kinetic components, but this is beyond the intent of the present evaluations.

### RKK mutants also increase likelihood that β2 subunits inactivate from closed states

BK β2 subunits, like β3a, also produce nearly complete inactivation of BK currents. However, in contrast to the β3a-mediated slow tails during recovery from inactivation, little or no channel openings are observed during recovery from β2-mediated inactivation (Solaro et al., 1997; Xia et al., 1999). Since inactivation in both cases presumably involves a similar access pathway to the central cavity (Zhang et al., 2006; Zhang et al., 2009), we therefore tested the impact of mutations of the RKK motif on β2-mediated inactivation.

Sample macroscopic currents of hβ2−containing WT, RKK3Q, RKK3E, and RKK3V channels before (left panels) and after (right panels) trypsin digestion are shown in Figure 13A,C,E,G along with corresponding GV curves at 0 and 10 μM Ca^2+^ following trypsin removal of β2-mediated inactivation (Fig. 13B,D,F,H). For a steady-state inactivation protocol, peak inactivating current of WT+hβ2 channels evoked by steps to +160 mV reaches approximately 0.70±0.07 (n = 5) of that after trypsin digestion. For RKK3Q+β2 channels, the peak of the inactivating current at +160 mV is 0.29±0.07 (n = 5) of that after trypsin treatment (Fig. 13C and E). For RKK3E+hβ2 (Fig. 13E) or RKK3V+hβ2 (Fig. 13G), the peak outward currents before trypsin digestion were 0.05±0.02 (n = 4) and 0.08±0.01 (n = 4) of that after trypsin digestion, respectively. Overall, these results parallel the steady-state inactivation of the D20Am-associated RKK mutant channels and are consistent with the idea that hβ2 subunits in association with RKK-mutated triplets are more likely to occupy inactivated states prior to the depolarizing voltage steps. Most importantly, the steady-state inactivation curves obtained with the RKK mutants with β2 (Fig. 13J) exhibit limiting fractional availability with no obvious voltage-dependence from −160 mV to −200 mV. As above, this can best be explained if there is an inactivated closed state in voltage-independent equilibrium with resting closed states. One other point regarding the β2-mediated effects is that, unlike D20Am, they are not associated with an increase in inward current at negative potentials. This absence of inward current almost certainly relates to the absence of channel openings during recovery from inactivation observed for Slo1+β2 currents (Xia et al., 1999) in contrast to the prolonged O*-I bursts associated with β3a(D20Am)-mediated inactivation (Zeng et al., 2007; Gonzalez-Perez et al., 2012). Thus, for the native β2 N-terminus, if there is a two-step inactivation process, something revealed in the FIW/FIE mutation of β2 (Benzinger et al., 2006), it may be the case that the state following N-terminus binding is also nonconducting.

**Figure 13.**
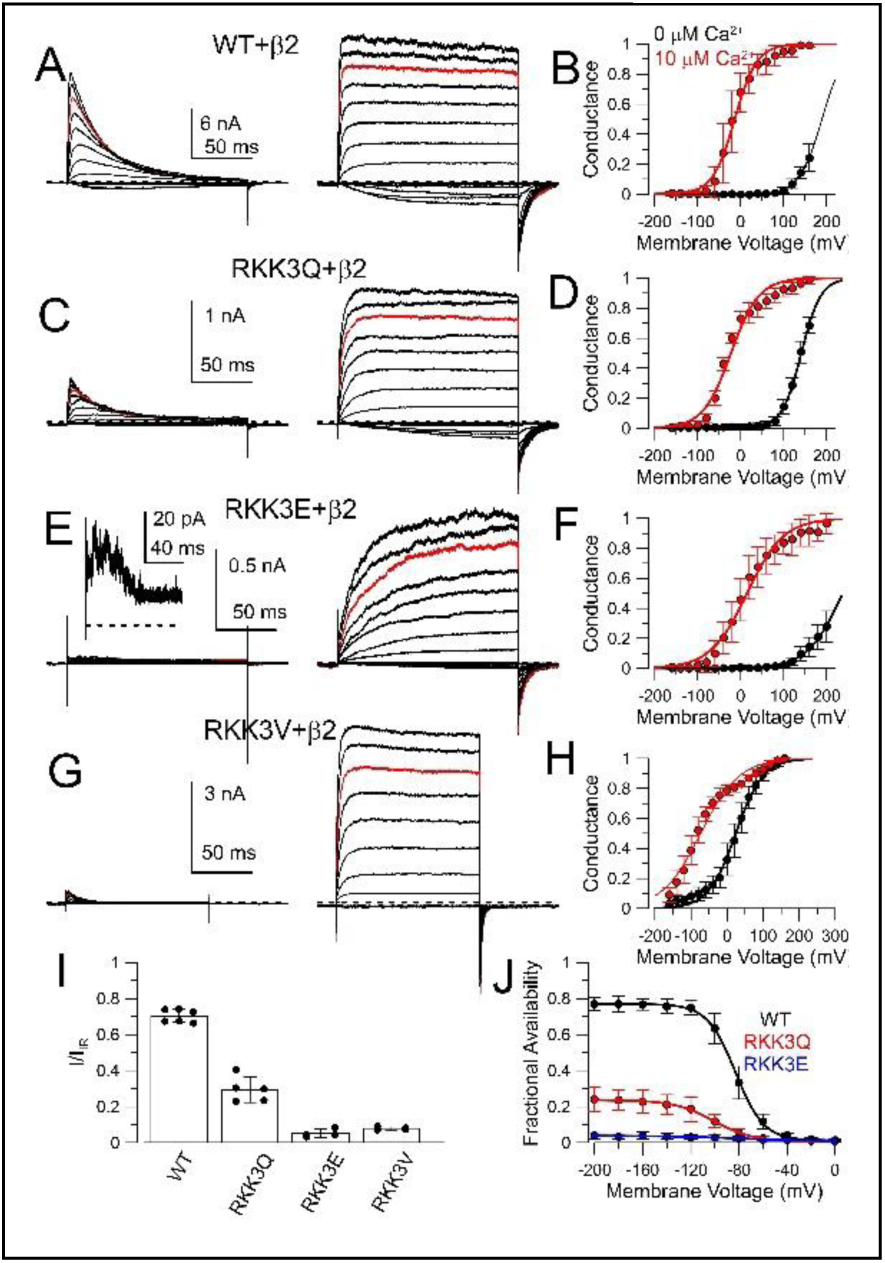
β2-associated RKK-mutant BK channels exhibit features of closed-state inactivation. **(A)** Currents resulting from Slo1 WT+β2 expression with 10 μM Ca^2+^ before (left) and after (right) trypsin-mediated removal of inactivation. **(B)** GV curves for Slo1 WT+β2-IR at 0 (black) and 10 (red) μM Ca^2+^. For WT+β2 (IR), at 0 Ca^2+^, V_50_= 188.8±13.2 mV, with z= 0.91±13.2 and, at 10 μM Ca^2+^, V_50_=-18.2±2.6 mV, with *z*=1.11±0.11*e* **(C)** Currents resulting from RKK3Q+β2 at 10 μM Ca^2+^, before (left) and after (right) trypsin**. (D)** GV curves at 0 and 10 μM Ca^2+^ for RKK3Q+β2-IR with, at 0 Ca^2+^, V_50_= 136.4±3.8 mV, with *z*= 1.19±0.20*e* and, at 10 μM Ca^2+^, V_50_=-31.6+4.2 mV, with *z*=1.02+0.15*e*. **(E)** RKK3E+β2 currents at 10 μM Ca^2+^, before (left) and after (right) trypsin. **(F)** GV curves at 0 and 10 μM Ca^2+^ for RKK3E+β2-IR. At 0 Ca^2+^, V_50_= 233.4±12.4 mV, with z= 0.58±0.12*e* and, at 10 μM Ca^2+^, V_50_=x2.2±3.2 mV, with *z*= 0.74±0.06*e*. **(G)** Currents resulting from RKK3V+β2 at 10 μM Ca^2+^, before (left) and after (center) trypsin. **(H)** GV curves at 0 and 10 μM Ca^2+^ for RKK3V+β2-IR. For 0 Ca^2+^, V_50_= 29.4±7.3 mV, with *z*= 0.70±0.13 and, at 10 μM Ca^2+^, V_50_=-79.0±7.4 mV, with *z*= 0.61±0.09*e*. **(I)** Mean and individual values of peak current amplitude for inactivating current normalized to maximal current after removal of inactivation for each patch. **(J)** Fractional availability curves for WT (n=5), RKK3Q (n=5), and RKK3E (n=4), each coexpressed with β2 (as in Fig. 4D). Fit parameters (mean±90% confidence limits) were: for WT, h_∞_=0.77±0.01; V_50_ = −82.4+0.8 mV, *z*=2.1±0.1e; for RKK3Q, h_∞_ =0.24±0.01; V_50_=-100.3±1.9 mV; z=1.4±0.1e; for RKK3E, h_∞_ =0.04±0.004; V_50_=-69.4±15.9 mV; z=0.49±0.1e. Comparison of fractional availability values at −200 mV yielded, for WT vs. RKK3Q, P<0.0001, for WT vs RKK3E, P<0.0001, and for RKK3Q vs. RKK3E, P<0.0001, with 2 way ANOVA with Tukey’s multiple comparison correction.

### E321/E324 mutations reduce features of D20Am-mediated inactivation

The RKK triplet has been proposed to stabilize closed conformations of the BK channel by interaction with acidic residues, E321 and E324, of adjacent S6 helices (Tian et al., 2019). Hypothesizing that some of the effects of the RKK mutant constructs might arise from disruption of this interaction, we therefore tested whether mutation of the E321/E324 pair might also mimic aspects of the RKK mutations. We expressed D20Am with BK subunits in which the E321/E324 motif was mutated to EE/QQ, EE/DD, EE/GG, EE/WW, and EE/KK. EE/QQ has been previously shown to result in decrease in single channel current (Brelidze et al., 2003), while also producing a leftward shift in gating (Tian et al., 2019). Macroscopic currents were generated before and then after removal of inactivation by trypsin (Fig. 14A1-E1). GV curves were determined at 0 and 10 μM Ca^2+^ for each construct (Fig. 14A2-E2) expressed alone (dotted lines) and then also for the construct expressed with D20Am but following trypsin digestion (solid symbols and fitted Boltzmann). For each construct+D20Am, time constants of inactivation (Fig. 14F) and activation following removal of inactivation (Fig. 14G) were determined. Finally, the maximal current with intact inactivation was normalized to that after removal of inactivation to ascertain whether there is increased likelihood that channels may occupy inactivated states prior to depolarization (Fig. 14H). For three constructs, EE/QQ, EE/GG, and EE/KK, the peak amplitude of the inactivating currents was greater than 0.9, suggesting that under any gating model there was likely to be little if any inactivation either prior to the depolarization or during the rising phase of current. Consistent with this, for each of these three constructs, it can be seen that they activate faster than the WT BK channels expressed with D20Am (Fig. 14G), while inactivation time constants are actually slowed relative to the WT situation (Fig. 14F). In contrast, for EE/DD and EE/WW, the peak current amplitude relative to maximal current following trypsin is similar to WT (Fig. 14H) and both constructs exhibit both slowed activation and slowed inactivation. In contrast to the RKK mutations, none of the E321/E324 mutations exhibit any indication of enhanced inactivation at negative potentials prior to depolarizing activation steps. For this set of EE mutations, there is no indication of a persistent inward current except for that expected based on the shift of the GV curve (e.g., Fig. 14C1 for EE/GG). It can also be seen that the EE mutations abolish the slow tails that are associated with WT D20Am inactivation, in marked distinction to the RKK mutations that enhance the slow tails. This would suggest that the consequences of the RKK mutants cannot be explained simply by disruption of any RKK/EE interaction. The absence of slow tails in the EE mutants may suggest binding affinity of the D20Am N-terminus within the central cavity is influenced by the EE motif. We have not yet tested simultaneous changes in the RKK and EE motifs.

**Figure 14.**
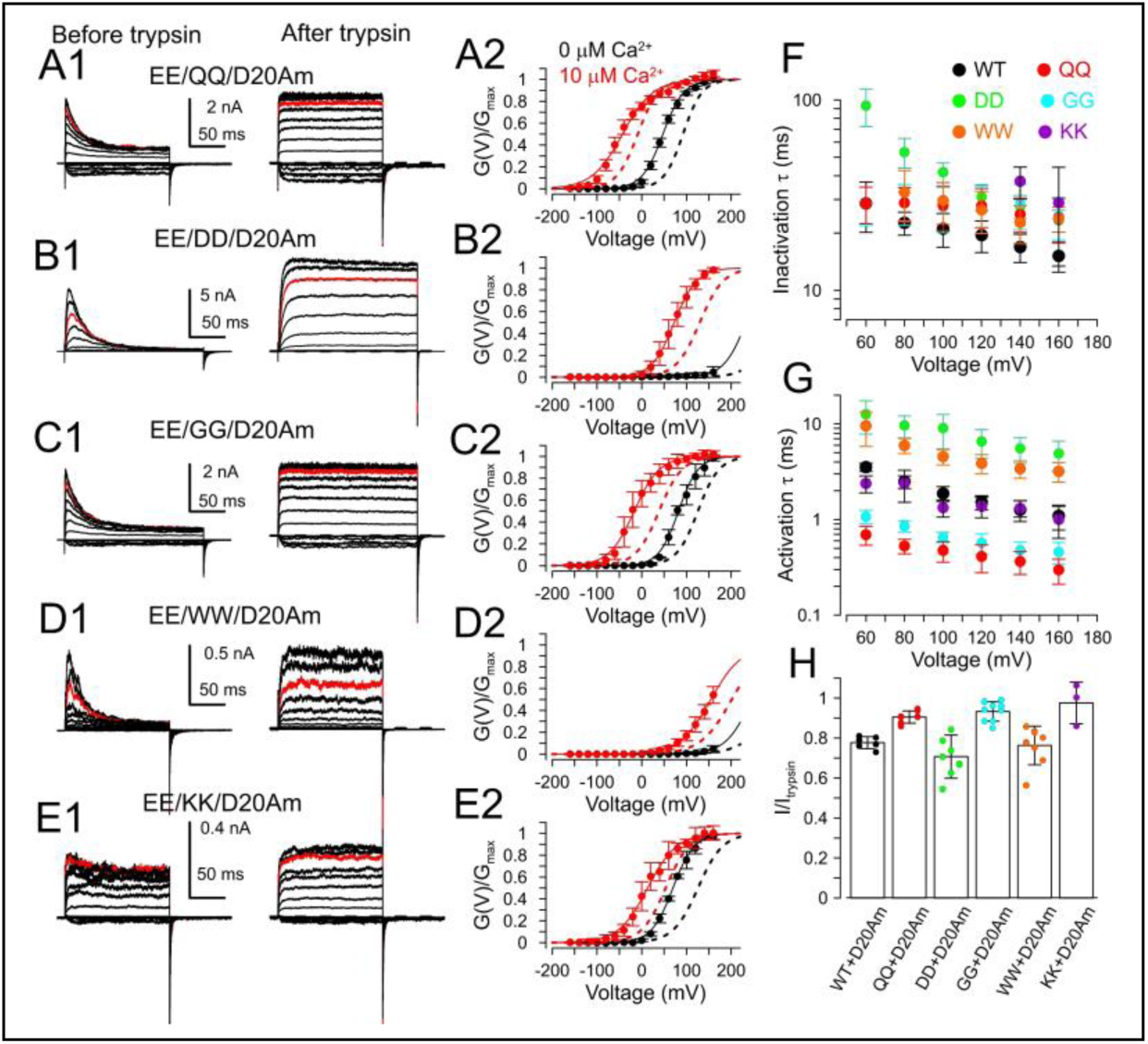
E321/E324 mutations do not result in persistent inward current or reduce likelihood of channel activation during depolarization, but do reduce or abolish slow tail current. (A1) A standard activation protocol with depolarization to +160 mV was applied to a mutated EE/QQ BK construct coexpressed with D20Am both before (left) and after (right) trypsin. (A2) GV curves from EE/QQ+D20Am-IR currents at 0 (black) and 10 (red) μM Ca^2+^. Dotted lines are EE/QQ GVs without D20Am subunits. Fitted Boltzmann values with D20Am-IR: for 0 Ca^2+^, V_50_= 47.3±3.0 mV, *z*=1.10±0.21*e*. For 10 μM Ca^2+^, V_50_ = −42.9±5.8 mV, z=0.75±0.11*e*. Without D20Am-IR, for 0 Ca^2+^, V_50_= 92.9±1.9 mV, *z*=1.20±0.09*e*. For 10 μM Ca^2+^, V_50_=-8.9±1.7 mV, *z*=1.09±0.07*e*. (B1) EE/DD+D20Am before and after trypsin. (B2) GVs for EE/DD without and with D20Am-IR. With D20Am-IR, for 0 Ca^2+^, V_50_= 233.2+10.5 mV, *z*=1.02+0.06*e* and, for 10 μM Ca^2+^, V_50_ = 72.7±1.8 mV, *z*=1.02±0.06*e*. Without D20AM-IR, for 0 Ca^2+^, V_50_= 293.8±14.4 mV, *z*=0.97±0.04*e* and, for 10 μM Ca^2+^, V_50_ = 129.6±1.7 mV, and *z*=0.97±0.04*e*. (C1) EE/GG+D20Am before and after trypsin. (C2) GVs for EE/GG without and with D20Am-IR. With D20Am-IR, for 0 Ca^2+^, V_50_= 82.2±4.7 mV, z=1.16±0.20*e* and, for 10 μM Ca^2+^, V_50_ = 16.7±4.0 mV, *z*=0.91±0.11*e*. Without D20AM-IR, for 0 Ca^2+^, V_50_= 125.6±3.1 mV, and *z*=1.08±0.11*e* and, for 10 μM Ca^2+^, V_50_ = 38.3±2.3 mV, *z*=0.99±0.07*e*. (D1) EE/WW+D20Am before and after trypsin. (D2) GVs for EE/WW without and with D20Am-IR. With D20Am-IR, for 0 Ca^2+^, V_50_= 245.5±21.3 mV, *z*=0.89±0.20*e* and, for 10 μM Ca^2+^, V_50_ = 154.5±0.53 mV, and *z*=0.73±0.01*e*. Without D20AM-IR, for 10 μM Ca^2+^, V_50_ = 201.2±52.0 mV, *z*=0.74±0.26*e*. The GV for EE/WW in 0 Ca^2+^ was not determined. (E1) EE/KK+D20Am before and after trypsin. (E2) GVs for EE/KK without and with D20Am-IR. With D20Am-IR, for 0 Ca^2+^, V_50_= 71.7±3.5 mV, *z*=1.05±0.12*e* and, for 10 μM Ca^2+^, V_50_ = 8.6±3.6 mV, and *z*=0.76±0.07*e*. Without D20AM-IR, for 0 Ca^2+^, V_50_= 125.7±3.4 mV, *z*=0.93±0.09*e* and, or 10 μM Ca^2+^, V_50_ = 53.4±3.4 mV, and *z*=0.96±0.09*e*. (F) Inactivation time constants for the indicated constructs, indicating that that all EE mutations slow the onset of inactivation. (G) Activation time constants measured after trypsin, showing that EE/QQ and EE/GG activate faster than WT, EE/WW and EE/DD somewhat slower, and EE/KK about the same as WT. (H) Peak inactivating outward current for each EE construct normalized to peak current after trypsin.

## Discussion

The Results in this paper are focused on trying to explain unusual phenomenology that arises from effects of mutations of the BK S6 RKK triplet on inactivation mediated by a β3a subunit N-terminal domain. Because of complexities in the Results, many of the implications of specific results and our attempts to explain them have already been discussed within the Results. Therefore, here we briefly summarize the main observations and the resulting conclusions. This is followed by a discussion of the implications of the results both for understanding the physical basis of BK inactivation and for consideration of closed-state inactivation in other channels.

### Summary of key observations

Compared to WT+D20Am current, coexpression of RKK mutant BK channels with D20Am reduces the peak inactivating outward current evoked by depolarizations, markedly slows tail currents, and results in appreciable persistent inward current at voltages for which the parent BK construct is largely closed. Both macroscopic and single channel results indicate that reduction in outward current arises from inactivation occurring at negative prepulses preceding the depolarizations, rather than during the rising phase of current activation. The persistent current at negative potentials arises from channels opening to a reduced conductance for which the fractional amplitude relative to normal openings increases with hyperpolarization, resulting in an apparently anomalous voltage-dependent increase in conductance as membrane potential is made more negative. The reduced conductance openings are consistent with effects of the isolated N-terminal β3a(1-20) L-peptide on single BK channels (Gonzalez-Perez et al., 2012) which has been explained as the result of rapid equilibrium between a preinactivated open state (O*) and an inactivated state (I). The voltage-dependence of the O*-I behavior is such that depolarization above 0 mV immediately favors channels in I, explaining why depolarization during the persistent current results in immediate inactivation, while, upon repolarization, the O*-I equilibrium favors instantaneous inward tail current reflecting greater fractional occupancy in O*.

In earlier work, in order to explain the large excess of net current flux during the macroscopic and single channel β3a tail current behavior over that predicted by recovery from a simple block model (I→O→C), the idea was proposed that β3a-inactivated BK channels recover from inactivation as follows: I←→O*→O→C (Zeng et al., 2007; Gonzalez-Perez et al., 2012). Building from that idea, here we extended the model describing D20Am+WT behavior to include activation transitions (Fig. 2C). This simple 5-state model was able to reasonably describe the major features of the reduction of outward current, prolongation of tail currents, and persistent tail (Fig. 10). Furthermore, many aspects of the differences in β3a-induced inactivation among WT and RKK mutants including reduction of outward current, tail current prolongation, and persistent inward current could be approximated by varying the O*→O rate (Fig. 11).

However, despite the utility of the linear 5-state model in Scheme 1, several aspects of the results suggest that, although the D20Am N-terminus is unlikely to reach a position of inactivation in closed WT channels, in the RKK mutated channels, the D20Am N-terminus can gain access to and bind to a position in the closed BK channel, resulting in a closed state inactivation. The evidence in support of this idea was three-fold. First, in steady-state inactivation experiments, D20Am-associated RKK3Q, RKK3E, and RKK3V all had voltage-independent ℎ_∞_ asymptotes which, after normalization to maximal BK conductance in a patch, were less than that for D20Am-associated WT channels. Second, in single channel experiments, we were able to directly identify a fraction of long duration closures at the end of the S1 prepulse hyperpolarization which led to a complete failure to open during the subsequent S2 depolarization. Third, evaluation of the steady-state level of tail currents following repolarization from strong, inactivating depolarizations revealed that a substantial fraction of RKK3Q and RKK3E channels that return to closed states during the tail currents remain unable to open upon depolarization. Together these results strongly support the conclusion that the β3a N-terminus can bind to resting closed states of the RKK-mutant BK channels in a position that prevents normal channel opening upon depolarization. We modeled this by inclusion of a C_1_←→C_1_* and C_2_←→C_2_* equilibria, which we infer is voltage-independent because of the voltage-independent ℎ_∞_ asymptote in the fractional availability curves.

Despite the lines of evidence supporting the existence of a long-lived inactivated state at negative potentials, it is interesting that occupancy in C_1_* and C_2_* states must be substantially reduced by the end of a 150 ms depolarizing inactivation step. This is supported by the fact that the single channel results show that following repolarization to −160 mV from +140 mV, almost all D20Am-associated channels almost always residues in O*-I at the time of repolarization. However, this apparent absence of channels in C_1_* or C_2_* at the end of depolarizations is easily explained by the idea that, during a depolarization, transitions from C_1_*→C_1_ and C_2_*→C2 are sufficiently rapid that channels are then able to open and transition into O*-I during the 150 ms depolarization prior to repolarization. A testable prediction of this idea is that, in single channel patches, following shorter inactivation steps during which C_1_*→C_1_ transitions may not yet have occurred, one might expect to see a larger fraction of repolarizations revealing not exhibiting any opening during tail currents, as channels simply transition from C_1_* to C_1_.

### Conceptualization of the onset of inactivation and the O*-I equilibrium

BK inactivation, whether mediated by β3a (Zeng et al., 2007; Gonzalez-Perez et al., 2012), β3b (Xia et al., 2000; Lingle et al., 2001), β3c (Uebele et al., 2000) or β2 (Wallner et al., 1999; Xia et al., 1999) subunits, involves a mobile cytosolic N-terminal peptide segment. For BK channel inactivation, many of the criteria used to argue that an N-terminus acts within the central cavity by simple channel block mechanism are not met, which has left some uncertainty regarding details of the mechanism of inactivation. For example, cytosolic blockers do not slow β-subunit mediated onset of inactivation (Xia et al., 1999; Xia et al., 2000), and tail currents during recovery from inactivation do not follow expectations for simple block behavior (Solaro and Lingle, 1992; Zeng et al., 2007). However, in contrast to simple block predictions, it has been shown that the 2-step inactivation model can explain both the absence of prolongation of inactivation onset by cytosolic blockers (Lingle et al., 2001) and also the β2- and β3a-mediated tail current behavior (Zeng et al., 2007). Moreover, although inactivation onset mediated by the β3a N-terminus is not hindered by TBA or bbTBA, both QA blockers do hasten the time course of deactivation of β3a-N-terminus mediated tail currents (Gonzalez-Perez et al., 2012), which is consistent with binding of either QA compound to the O* state, while hindering formation of the I state. This supported the idea that the I state involves occupancy of the N-terminus in a position that can also be hindered by the presence of TBA or bbTBA, which one presumes is near the selectivity filter.

How might the O*-I behavior be explained? We would envisage that, for the β3a N-terminus, O* represents binding of the N-terminus in a position within the central cavity which still permits ion permeation, but that mobility of that N-terminus or some element of the N-terminus, perhaps as simple as a single side chain, produces rapid transient occlusion of outward current flux, while inward current flux more strongly favors occupancy in O*. It should be noted that the first 15 residues of the β3a N-terminus are entirely uncharged (Fig. 2). Two topics for future consideration are: 1. whether ion occupancy in the permeation pathway may determine or impact on the voltage-dependence (z=∼0.25*e*) of β3a-mediated inactivation (which largely arises from its first 15 uncharged N-terminal residues (Gonzalez-Perez et al., 2012) and 2. determination of the dependence of O*-I behavior on specific residues in the β3a N-terminus and BK S6.

### Conceptualization of closed-state inactivation

A significant new point arising from the present work is that, at least in the mutant RKK channels, the D20Am N-terminus can access a binding site in closed channels. The steady-state inactivation curves, the observation of a long duration closed-state at negative potentials preceding depolarizations that fail to result in openings of channels, and then that a fraction of channels that return to closed states in tail currents are unable to be activated all support this idea. Although inactivation from closed states has often been used as a component of kinetic models that may improve the ability of a model to account for relative positions of activation GVs and steady-state availability curves along the voltage-axis (Zagotta and Aldrich, 1990; Ayer and Sigworth, 1997), direct evidence for closed-state inactivation can be difficult to obtain. In the present case, three lines of evidence point directly to the existence of a closed, inactivated state in equilibrium with a resting closed state. We suggest that this reflects a voltage-independent C_1_←→C_1_* equilibrium, reflecting binding of the β3a N-terminus within the central cavity while the RKK-mutated channels are in the most resting closed state. The requirement that this must be voltage-independent, which arises from the voltage-independence of the ℎ_∞_ asymptotes, is the basis for our use of a voltage-dependent O←→O* equilibrium in simulations.

Functionally, we envision this as follows. In the RKK-mutant channels, the D20Am N-terminus can enter the central cavity via the disrupted side portal region of S6 reaching a binding site that is essentially equivalent to that reached by the N-terminus in open channels. We would suggest that whether the channel is occupied by the N-terminus in a closed conformation or an open conformation, transition between C*←→O* is hindered by the presence of the N-terminus. This suggestion arises, since otherwise repolarization when channels are in the O*-I ensemble would rapidly lead to channels returning to closed states, preventing the occurrence of slow tail currents. Given that there are faster components in the experimentally observed tail currents than are seen in the simple simulations, it may be possible that C*←→O* transitions can occur, albeit at rates much slower than the C←→O transitions. This will require additional evaluation within the context of a more realistic BK gating scheme.

Our results do not clearly support the idea that resting closed-state inactivation can occur with wild-type channels, although we cannot fully exclude that possibility. Potential closed-state inactivation might be considered of two types: 1) inactivation analogous to what we observe for the RKK-mutated BK channels in which the most resting closed state can become occupied by the N-terminus or 2) that inactivation of a closed state can only occur in partially activated closed states, in which some preopening conformational change permits the N-terminus to reach its binding site. In the present results, it is possible that closed-state inactivation of the second type, i.e.,from states leading to activation, may contribute to the leftward shift of WT steady-state inactivation curves relative to the simulated ℎ_∞_ curve. This ability of closed-state inactivation to shift ℎ_∞_ curves leftward has also been a consideration in early models of Shaker Kv channels (Zagotta and Aldrich, 1990; Ayer and Sigworth, 1997). For BK channels, closed state inactivation has been previously implicated in results with patches of single β2-mediated inactivation in rat chromaffin cells with prepulses to −40 mV at low Ca^2+^ resulting in inactivation of BK channels without any detectable opening, whereas direct steps from −120 reliably elicited openings at +60 mV (Ding and Lingle, 2002). Perhaps reflecting changes in pore access among closed states, although BK channel inhibition by bbTBA was shown to occur when channels are closed (Wilkens and Aldrich, 2006), subsequent work presented evidence that bbTBA did not block BK channels under conditions that voltage-sensors may be completely inactive, but that block could occur prior to channel opening (Tang et al., 2009). Overall, all results point to the idea that, in fully resting closed states, WT BK channels have barriers that prevent access of β subunit N-termini and bulky cytosolic blockers to positions of occlusion. However, conformational changes that occur that precede opening seem to be permissive for both β subunit mediated-inactivation and blockade by bulky QA compounds.

### Might resting closed state inactivation be a generally overlooked phenomenon*?*

The generation of steady-state inactivation curves is a standard tool used in the description of inactivating currents. Typically, the SSI curves are normalized to the peak current activated by the strongest depolarization in the protocol from the most negative conditioning potential, such that the ℎ_∞_ asymptote is 1.0. In the present work, taking advantage of the ability of trypsin to rapidly remove BK inactivation, SSI curves were normalized to the actual maximal BK conductance in a patch, thereby revealing that the limiting fractional inactivation varied dependent on the RKK mutation being studied. Typically, it is not possible or it is simply very difficult to undertake a normalization to absolute conductance. To our knowledge, it has never been attempted for voltage-dependent Na^+^ (Nav) channels, although, in a study of the L382I mutation in Shaker channels (Ayer and Sigworth, 1997), removal of inactivation by trypsin was used to argue that there was an inactivation pathway in this mutant that reduced the number of available channels for activation. An implication of the present results is that the ability to define absolute fractional availability for activation via normalization to maximum conductance in a patch or cell can potentially provide insight into how closed state inactivation might be altered whether by mutations or by physiological regulatory processes.

### Comparison of D20Am binding to open and closed states

From the perspective of the 2-step inactivation formulation, the equilibrium constant for N-terminus binding within the BK central cavity is given by O*/O. Whereas the rate of O→O* is guided by the macroscopic inactivation time constant, O*→O can only be indirectly inferred via our model-dependent simulations. The values used to approximate WT inactivation behavior led to K_eq_=60/70. In our simulations that approximate the effects of RKK mutations, we assumed that k_3_ was unaffected by the mutation, while only k_-3_ was altered. The ℎ_∞_ values defined for RKK3Q, RKK3E, and RKK3V, following correction for the approximately 10-15% of channels that may inactivate prior to peak outward current were 0.44, 0.10, and ∼0.02, respectively. Interestingly, assuming that the relative rates of entry and unbinding from closed states are similar to those for open state inactivation, the predicted ℎ_∞_ values for our simulations with k_3_/k_-3_ ratios of 60/30, 60/10, and 60/2 are 0.33, 0.14, and 0.03, not too dissimilar from the experimental measurements for 3Q, 3E, and 3V. These values would require that the equilibrium constant for closed state occupancy is remarkably similar to that for open state occupancy, suggesting the binding site that mediates stereospecific binding of the β3a N-terminus is not dramatically altered between open and closed channels. Given that the β3a(1-20) peptide mimics most of the unique functional effects of the tethered β3a N-terminus, timed peptide applications to closed or open channels may provide a way to assess directly closed and open state affinities.

### Destabilization of EE/RKK interactions do not seem to underlie the enhanced β3a and β2 mediated inactivation

Interactions between E321/E324 and the RKK triplet were proposed to contribute to stabilization of the channel closed state (Tian et al., 2019). The present results indicate that E321/E324 mutations do not mimic the RKK effects on inactivation, but, rather, EE mutations produce effects qualitatively consistent with a weakened interaction between the β3a inactivation domain and its binding site. Thus, destabilization of the EE/RKK interaction is not the basis for the enhanced stability of β3a or β2-mediated inactivation. However, both RKK and EE do impact on the stability of the inactivated states, with RKK mutations enhancing occupancy of inactivated states and EE mutations weakening the stability of inactivated states. We suggest that the specific identity of residues in the RKK triplet either directly or indirectly stabilizes binding of the β N-termini within the central cavity, most dramatically illustrated by RKK3V which essentially locks channels in O*-I states at least under the duration of protocols used in our experiments. Although it is tempting to also suggest the E321/E324 may also stabilize binding of the β3a N-terminus within the central cavity, mutation of the only three basic residues in the first 20 residues of β3a (R16QR17QR18Q) does not prevent the O*-I tail current behavior (Gonzalez-Perez et al., 2012). Although it is natural to try to interpret changes in β3a-mediated inactivation in terms of the residues altered in the RKK or EE mutations, it is important to keep in mind that there is also increasing evidence that the central cavity of BK channels and particularly in the vicinity of the RKK motif is likely decorated with lipids (Kallure et al., 2023), which may be altered by the RKK or EE mutations.

### Can O*-I behavior be explained by a single O* state, i.e., a single open state with marked current rectification?

One point that occasionally arises in regards to consideration of the β3a tail current phenomenology is whether the O*-I proposal might be approximated just as well by a single open state with marked current rectification. There are several arguments against this possibility. First, the voltage-dependence of the O*-I current amplitude relative to the O state amplitude fits very well with a simple two-state blocking process. Other cases for which channels exhibit such marked current rectification often arise because of known blocking particles, such as Mg (Davies et al., 1996) or spermine (Lopatin et al., 1994; Fakler et al., 1995), which are also best approximated by a two state equilibrium. Second, cytosolic quaternary ammonium blockers of BK channels do not slow the time course of development of β2 or β3-mediated inactivation (Solaro et al., 1997; Xia et al., 2000; Li et al., 2007). A one-step inactivation process, whether it involves or rectifying conductance or not, will be slowed by such blockers (Choi et al., 1991), whereas two-step inactivation in which the O→O* step is rate limiting, but blocker can bind to both states, will not exhibit QA-dependent slowing of inactivation (Lingle et al., 2001). On balance, the idea that binding of the β3a subunit N-terminus within the BK central cavity is a process separate from that producing channel occlusion remains the better explanation of the available data. The challenge ahead is to define determinants of β3a binding and those involved in the physical basis of rapid occlusion.

## Abbreviations used in this paper

BK, large conductance Ca^2+^-activated K^+^ channel

Po, open probability

GV, conductance/voltage

Kv, voltage-dependent K^+^ channel

TEA, tetraethylammonium

V_50_, voltage of half activation

G_max_, maximal conductance

ℎ_∞_, h-infinity, fractional availability of channels for activation.

## Author contributions

Y.Z. designed research, performed experiments, analyzed data, and prepared the manuscript. X.-M.X. contributed new reagents/analytical tools. C.J.L. designed research, analyzed data, and prepared the manuscript.

## Acknowledgements

This work was supported by R35GM118114 to CJL. We thank Vivian Gonzalez-Perez and Sandipan Chowdhury for encouragement and critical comments on the manuscript. Any supporting materials are available from the authors upon reasonable request.

## Supplementary Information

**Table S1.** GV curve parameters from WT and RKK mutant constructs.

| | 0 $\mu\text{M}$ $\text{Ca}^{2+}$ | | 10 $\mu\text{M}$ $\text{Ca}^{2+}$ | |
| --- | --- | --- | --- | --- |
| | $V_{50}$ (mV) | $z$ (e) | $V_{50}$ (mV) | $z$ (e) |
| WT | $178.1 \pm 5.4$ | $0.94 \pm 0.15$ | $30.9 \pm 1.7$ | $1.24 \pm 0.09$ |
| RKK3Q | $236.9 \pm 10.1$<br>$223.5 \pm 21.6^*$ | $0.89 \pm 0.18$<br>$0.86 \pm 0.17$ (Q)* | $90.3 \pm 1.5$ | $1.22 \pm 0.07$ |
| RKK3E | $282.9 \pm 16.4$<br>$265.3 \pm 30.8^*$ | $0.64 \pm 0.11$ | $135.4 \pm 1.2$ | $1.03 \pm 0.04$ |
| RKK3V | $38.9 \pm 6.5$<br>$29.5 \pm 28.0^*$ | $0.49 \pm 0.06$<br>$0.48 \pm 0.12^*$ | $-26 \pm 6.9$ | $0.42 \pm 0.04$ |

\*(Tian et al., 2019)

### Gating shifts produced by D20Am are unlikely to account for direct entry into O*-I states at negative potentials

To probe properties of inactivation arising from coexpression of D20Am with RKK mutant constructs, we first examined activation of RKK mutants+D20Am currents in order to determine conditions for which prepulses would favor occupancy of channels in closed states prior to test depolarizations. Based on Table S1, the RKK mutations themselves do not cause major leftward shifts in channel activation. Since inactivation precludes accurate determination of GV curves, for WT and RKK mutant subunits we measured currents following removal of inactivation by trypsin both at 0 and 10 μM Ca^2+^ (Fig. S1A-D1). From such records, tail currents were used to generate GV curves which were fit with single Boltzmann functions (Fig. S1A2-D2). The GV curves reveal that, for WT, RKK3Q, and RKK3E constructs when associated with D20Am-IR, even at 10 μM Ca^2+^, open probability is low at voltages negative to −100 mV, while maximal outward current activation is evoked at depolarizations of +160 mV. In contrast, the RKK3V mutation shifts gating substantially leftward, with or without D20Am-IR, such that the Po with 0 Ca^2+^ may be around 0.05 at −100 mV, although decreasing appreciably by −160 mV. Thus, some low opening probability might be expected at 0 Ca^2+^ for RKK3V at −160 mV. To confirm the general inferences made from the macroscopic GV curves regarding channel Po at negative potentials at a given Ca^2+^, we measured single channel activity for the two most left-shifted constructs, RKK3Q and RKK3V in association with D20Am-IR (Fig. S1E,F). This confirmed that RKK3Q+D20Am-IR channels rarely open at −160 mV with 10 μM Ca^2+^, while RKK3V+D20Am-IR channels rarely open at −160 mV with 0 μM Ca^2+^. However, we note that in some RKK3V+D20AM-IR patches, occasional openings were observed at −160 mV with 0 Ca^2+^ (Fig. 5D). However, overall, the occurrence of persistent current at negative potentials in the RKK mutant constructs when expressed with D20Am-IR is unlikely to result from a pronounced shift in the C-O equilibrium towards negative potentials, unless it arises from some unexpected effect of the D20Am N-terminus that is removed by trypsin.

**Figure S1.**
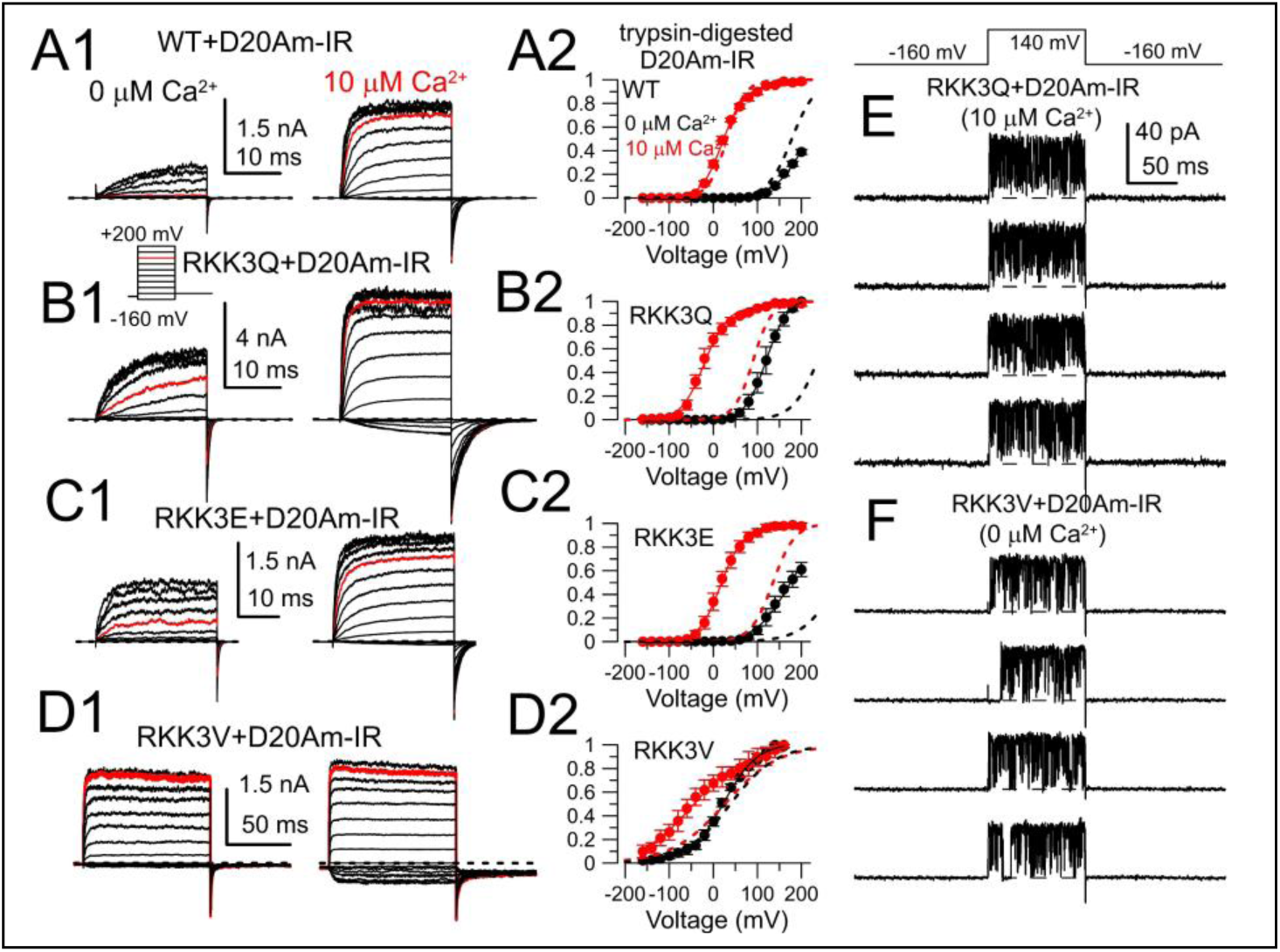
Activation curves of RKK-mutated subunits when coassembled with inactivation-removed D20Am subunits. **(A1)** Currents of WT channels in excised inside-out patches were activated by the voltage protocol shown in the inset at 0 (left) or 10 μM (right) Ca^2+^. Red traces: +120 mV. Patches were held at −140 mV for 50 ms prior to each depolarizing step**. (A2)** GV curves from WT+D20Am-IR channels. Dotted lines correspond to GVs for WT currents without coexpression with D20Am (values in Table 1). The Boltzmann fits are: V_50_ = 211.1 ± 5.1 mV, *z* = 0.71 ± 0.09*e* (0 Ca^2+^); V_50_ = 22.1 ± 2.1 mV, z = 1.01 ± 0.07*e* (10 μM Ca^2+^). (**B1**) Currents from RKK3Q BK channels for 0 and 10 μM Ca^2+^. (**B2** GV curves for RKK3Q+D20Am-IR channels, with dotted lines for RKK3Q alone: : V_50_ = 120.7 ± 4.1 mV, *z* = 1.04 ± 0.13*e* (0 Ca^2+^); V_50_ = −20.2 ± 3.1 mV, *z* = 0.96 ± 0.10*e* (10 μM Ca^2+^). (**C1**) Currents for RKK3E BK channels. (**C2**) GV curves for RKK3E+D20Am-IR: V_50_ = 172.2 ± 3.4 mV, *z* = 0.65 ± 0.06*e* (0 Ca^2+^); V_50_ = 17.1 ± 2.4 mV, *z* = 1.00 ± 0.08*e* (10 μM Ca^2+^). (**D1**) Currents for RKK3V BK channels. (**D2**) GV curves for RKK3V+D20Am-IR: V_50_ = 23.8 ± 6.2 mV, *z* = 0.64 ± 0.10*e* (0 Ca^2+^); V_50_ = −47.0 ± 8.5 mV, *z* = 0.44 ± 0.06*e* (10 μM Ca^2+^). (**E**) Openings of a single RKK3Q+D20Am-IR channel in 10 μM Ca^2+^, showing reduced Po at −160 mV. (**F**) Openings of a single RKK3V+D20Am-IR channel at 0 μM Ca^2+^, also exhibiting low Po at −160 mV.

**Figure S2.**
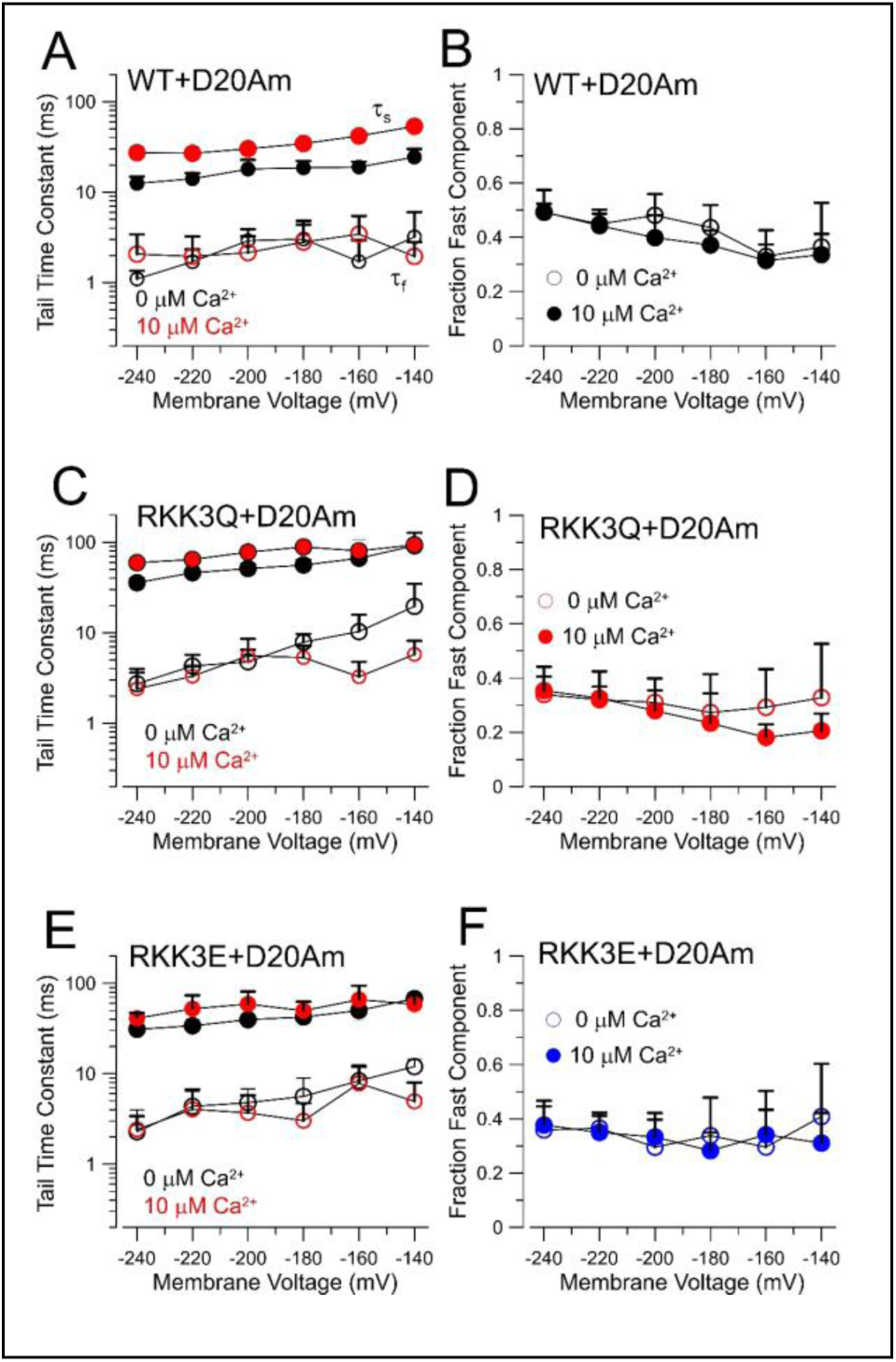
Time constants of deactivation for WT, RKK3Q, and RKK3E when expressed with D20Am. Tail currents were fit with a two exponential function as shown in Figure 9 for currents activated by either 0 or 10 μM Ca^2+^. Time constants and fractional amplitude of the fast component are plotted in Fig. S2A,B for WT+D20Am, in Fig. S2C,D for RKK3Q+D20Am, and in Fig. S2E,F for RKK3E+D20Am. Overall, time constants are faster with hyperpolarization. The slow time constant is somewhat slowed by increased Ca^2+^, while the fractional amplitude of fast and slow components does not seem appreciably altered by either voltage of Ca^2+^. **(A)** Time constants of fast (open symbols) and slow (filled symbols) for 0 (blue) and 10 (red) μM Ca^2+^ are plotted over voltages from −140 to −240 mV. **(B)** Fraction of the fast component of tail current for WT tail currents at 0 (open symbols) and 10 (closed symbols) μM Ca^2+^. **(C)** Time constants of fast and slow tail current components for RKK3Q+D20 Am. **(D)** Fraction of fast tail component for RKK3Q+D20Am. **(E)** Time constants of fast and slow tail current components for RKK3E+D20Am. **(F)** Fraction of fast tail component for RKK3E+D20Am at 0 and 10 μM Ca^2+^.

